# Controlling Neocortical Epileptic Seizures using Forced Temporal Spike-Time Stimulation: An *In Silico* Computational Study

**DOI:** 10.1101/2022.10.09.511502

**Authors:** Joseph Schmalz, Rachel V. Quinarez, Mayuresh V. Kothare, Gautam Kumar

## Abstract

Epileptic seizure is typically characterized by highly synchronized episodes of neural activity. Existing stimulation therapies focus purely on suppressing the pathologically synchronized neuronal firing patterns during the ictal (seizure) period. While these strategies are effective in suppressing seizures when they occur, they fail to prevent the re-emergence of seizures once the stimulation is turned off. Previously, we developed a novel neurostimulation motif, which we refer to as “Forced Temporal Spike-Time Stimulation” (FTSTS) [1] that has shown remarkable promise in long-lasting desynchronization of excessively synchronized neuronal firing patterns by harnessing synaptic plasticity. In this paper, we build upon this prior work [1] by optimizing the parameters of the FTSTS protocol in order to efficiently desynchronize the pathologically synchronous neuronal firing patterns that occur during epileptic seizures using a recently published computational model of neocortical-onset seizures [2]. We show that the FTSTS protocol applied during the ictal period can modify the excitatory-to-inhibitory synaptic weight in order to effectively desynchronize the pathological neuronal firing patterns even after the ictal period. Our investigation opens the door to a possible new neurostimulation therapy for epilepsy.

## 1 Introduction

Epilepsy affects 65 million people world wide and is typically characterized by highly synchronized episodes of neural activity that can lead to the loss of autonomy [3, 4]. In most cases, epileptic symptoms are treatable with anti-epileptic drugs, although not all patients respond to the drugs [5–7]. This type of epilepsy is coined as drug resistant epilepsy (DRE). Patients with DRE become prime candidate for direct neurostimulation therapies, such as vagus nerve stimulation [8, 9] or deep brain stimulation (DBS) [10], which have been proven to be effective in reducing the epileptic episodes. In case of DBS based therapy, the electrodes are implanted into a specific part of the brain prone to initiate epileptic seizure and high frequency electrical stimulation is applied to suppress the seizure activity [11]. While this approach works well at suppressing the seizure when it arises, it doesn’t address the reversal of the underlying network dynamics that generates seizures. As a result, seizures reemerge once the DBS is turned off and require its continued re-application.

Recently, we developed a novel neurostimulation strategy called “Forced Temporal Spike-Time Stimulation” (FTSTS) which showed, in simulation, promising results in efficiently desynchronizing large excitatory-inhibitory (E-I) spiking neuron networks by harnessing the long-term synaptic plasticity and keeping the network in the desynchronized state over a long time horizon after the stimulation was turned off [1]. Our FTSTS strategy consists of two biphasic out-of-phase pulses. One of the pulses is delivered to the excitatory neuron population and the other to the inhibitory neuron population. The pulse pair controls the relative spike times of each neuron population in order to control the average synaptic weight of the network and moves the E-I network from the synchronous to the asynchronous state. Furthermore, if the pulse-pairs are exactly reversed, the FTSTS protocol is capable of synchronizing an asynchronous neuron population. In simulation, we showed the capability of this selective FTSTS protocol in desynchronizing and resynchronizing neuron activity in a generic excitatory-inhibitory (E-I) network model of various sizes and dynamics. Based on these promising results, we wondered whether this FTSTS strategy could potentially be applied to terminate epileptic seizures effectively.

To investigate this question, in this paper, we consider a recently published biophysically constrained *in silico* model of a neocortical-onset seizure which has been validated using seizure data from human epileptic patients [2]. The model captures the key features of the underlying neocortical-onset seizure dynamics, such as the fast inward moving ictal discharges and the slow outward wavefront of ictal recruitments. Additionally, the model captures the predisposition of subsequent seizures after an initial seizure. Using this model, we systematically explore the parameter space of our FTSTS protocol to investigate the efficacy of the FTSTS strategy in reducing or increasing the neocortical-onset seizure prevalence. To determine the efficacy of the FTSTS protocol, we measure the changes in the average excitatory-to-inhibitory synaptic weights over the period of the applied FTSTS protocol.

In order to overcome the selectivity constraints inherent in the electrical FTSTS protocol as it targets excitatory and inhibitory populations, we further extend our electrical FTSTS protocol to a realistic and realizable selective optogenetic FTSTS protocol by integrating two optogenetic channelrhodopsin dynamic models i.e., Chronos and Chrimson, into the neocortical-onset seizure model.

## 2 Results

### Novel FTSTS Reduces Seizure Prevalence

We begin this section by showing simulation results from the *in silico* seizure model [2] described in Section 4 to illustrate the changes in the network synaptic strengths mediated by the episodes of seizure. We initiated a seizure in the model using a seizure initiating input of 200 pA applied for 3 sec to 50 neocortical excitatory neurons in the network of 500 excitatory (E) and 500 inhibitory (I) neurons. Figure 1A shows the raster plot of the emergence of three episodes of seizure.

**Figure 1:**
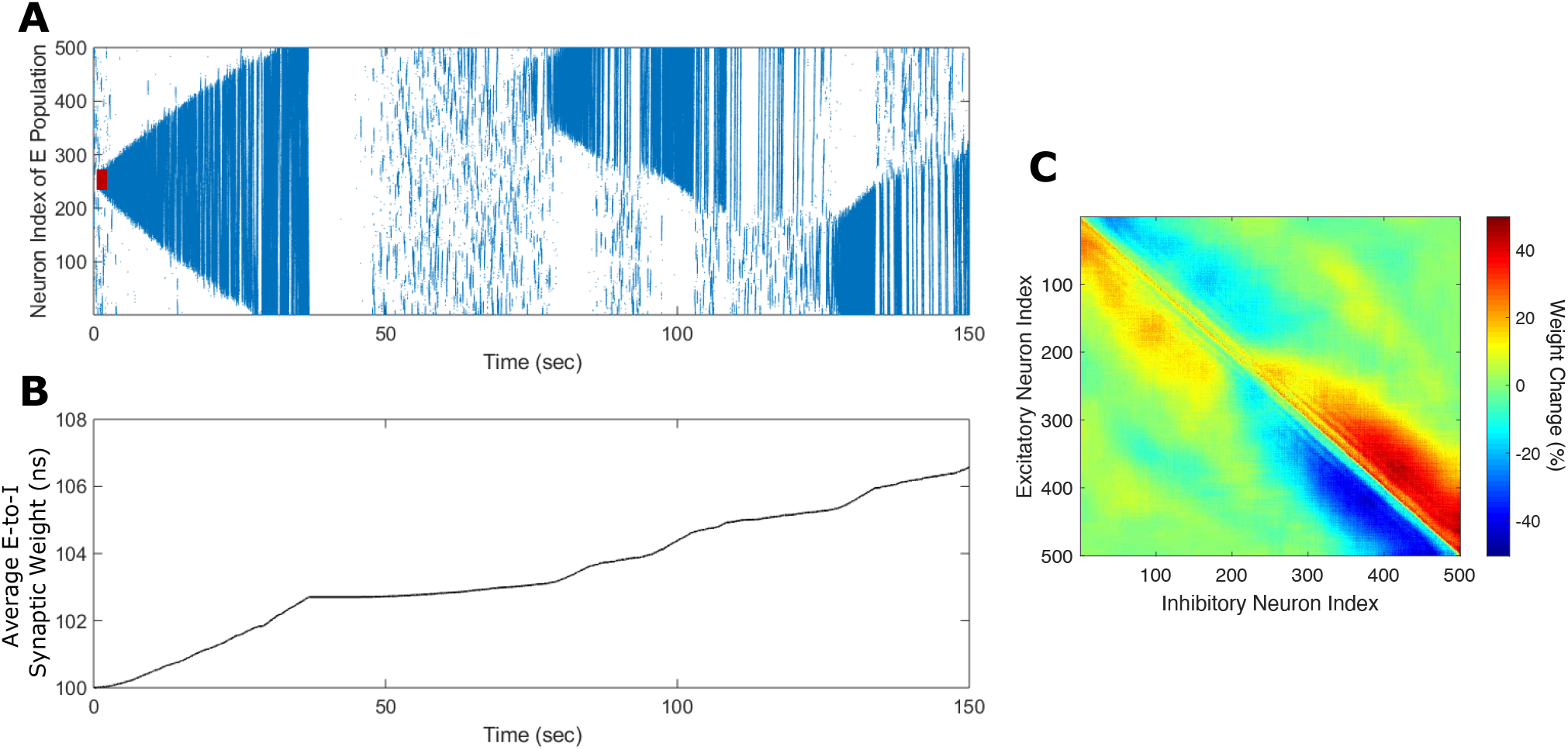
Propagation of the neocortical seizure through network. A seizure initiating input applied for 3 *s* (red-bar) to the *in silico* seizure model from [2] initiates a seizure. **(A)** shows the propagation of the initial seizure and the emergence of a second seizure after the first seizure terminates. **(B)** shows the increase in the average excitatory to inhibitory (E-to-I) synaptic weight during each seizure event. **(C)** shows the percent change in the strength of E-to-I synaptic weight of each synapse at the end of the simulation (*t* = 150 sec).

The first episode of seizure propagated through the entire network and terminated after 40 sec. After the initial episode of seizure, spontaneous spiking activity was observed for approximately 50 sec. Then, a second seizure spontaneously emerged, which was followed by a third seizure. As the seizure propagated throughout the network, the plastic synaptic connections between the excitatory and inhibitory (E-to-I) neurons were rewired by the seizure-like activity. In particular, the average E-to-I synaptic weight of the network increased during and after each seizure episode (see Figure 1B). More importantly, the synaptic weights were significantly modified by the seizure such that synaptic projections from the excitatory to the inhibitory neurons in front of the seizure wave were weakened and, conversely, those behind the seizure wave front were strengthened (see Figure 1C). This patterned rewiring prevented the inhibitory neurons in front of the seizure from inhibiting the seizure progression through the network. Next, we applied our FTSTS protocol to this *in silico* seizure model to investigate whether our protocol could suppress the prevalence of spontaneous seizures (see Section 4 for the details of the FTSTS protocol). After initiating a seizure in the model by applying the seizure initiating input (200 pA applied for 3 sec to 50 neocortical excitatory neurons), we applied the inverted-standard FTSTS protocol (see Figure 2D and Figure 13C) at the 10 sec mark for 5 sec (see Table 1 for the FTSTS parameters). The inverted-standard FTSTS protocol consists of the excitatory population pulses having the polarity of −1 (i.e., the positive phase of the biphasic pulse followed by the negative phase) and the inhibitory population pulses having the polarity of 1 (i.e., the negative phase of the biphasic pulse followed by the positive phase). In this setting, the applied FTSTS protocol forces more neurons to fire in a temporal pattern of pre-synaptic (excitatory) neurons before post-synaptic (inhibitory) neurons. Thus, the stimulation protocol leads to increase in the average E-to-I synaptic weight of the network and reduces the chances of the spontaneous emergence of future episodes of seizure.

**Figure 2:**
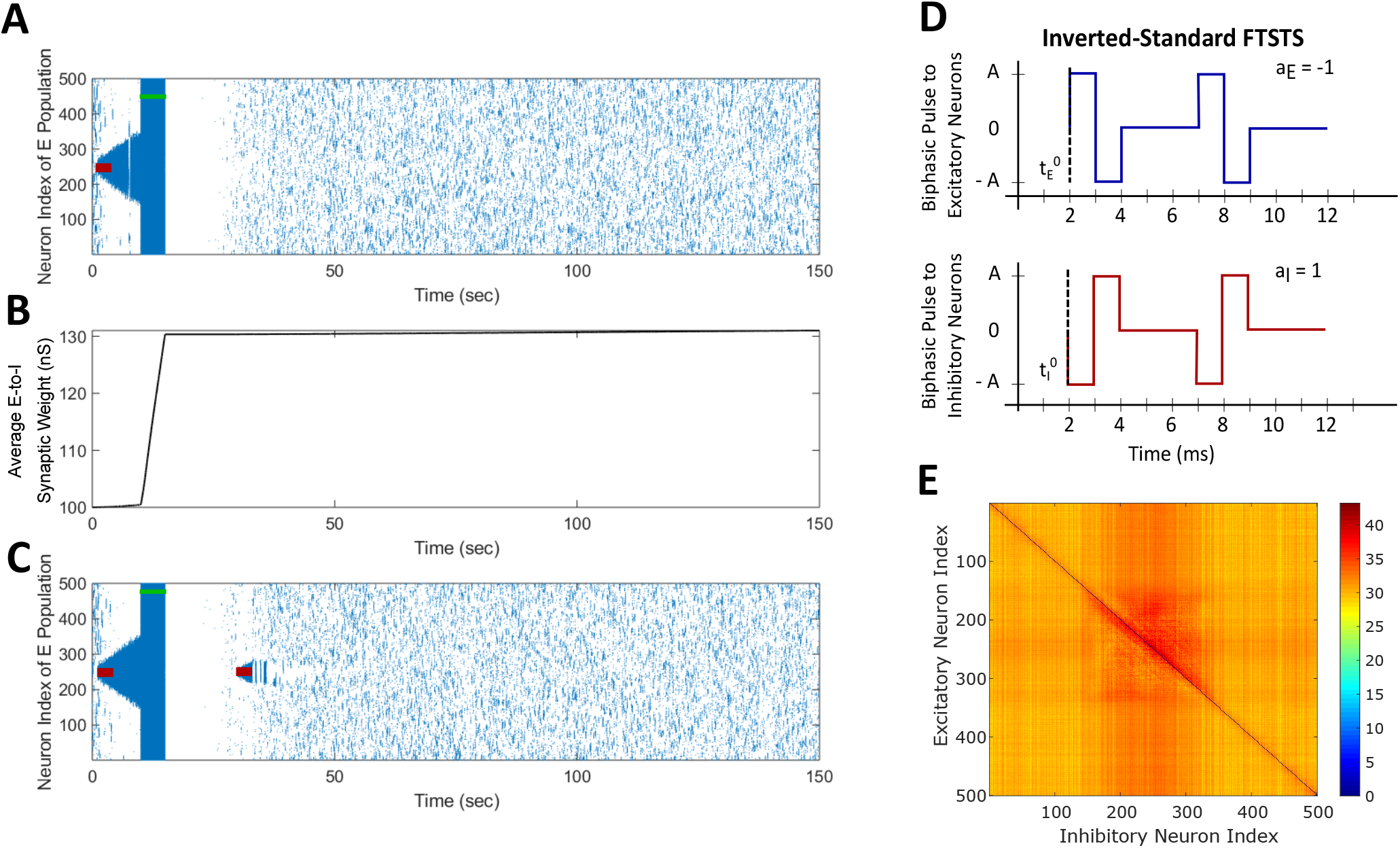
Suppression of spontaneous seizure with FTSTS. A seizure initiating input applied for 3 sec (red-bar) to the *in silico* seizure model initiated a seizure that begins to propagate through the network. **(A)** shows the suppression of neocortical seizure activity by our inverted-standard FTSTS protocol (green-bar). Our FTSTS protocol increased the average synaptic weight **(B)** while it was applied. Then, we applied a second seizure input for 3 sec at 30 sec. **(C)** shows that during the application of the second seizure initiating input a new seizure starts to propagate. As soon as the second seizure input is turned off, then the seizure stops propagating. **(D)** shows the inverted-standard FTSTS stimulation protocol where the polarity of the biphasic pulse delivered to the excitatory population is *a*_*E*_ = −1 and the polarity of the biphasic pulse delivered to the inhibitory population is *a*_*I*_ = 1. **(E)** shows the percent change in the excitatory-to-inhibitory synaptic weights at the end of the simulation (*t* = 150 sec in **(A)**).

**Table 1:** FTSTS Protocols and Parameters.

| FTSTS Parameters | Inverted-Standard | Standard | Mirrored | Inverted-Mirrored |
| --- | --- | --- | --- | --- |
| $a_E$ | -1 | 1 | 1 | -1 |
| $a_I$ | 1 | -1 | 1 | -1 |
| Pulse Waveform | bi-phasic square | bi-phasic square | bi-phasic square | bi-phasic square |
| Amplitude (nA) | 2 | 2 | 2 | 2 |
| Frequency (Hz) | 83 | 83 | 83 | 83 |
| Pulse Width (ms) | 1 | 1 | 1 | 1 |
| Interpulse Interval (ms) | 10 | 10 | 10 | 10 |
| Train-Offset Time (ms) | 0 | 0 | 0 | 0 |
| Duration (sec) | 5 | 5 | 5 | 5 |

As shown in Figure 2A, the inverted-standard FTSTS protocol disrupted the synchronous spiking activity of neurons and terminated not only the initiated seizure immediately but also the future episodes of spontaneous seizure activity. Furthermore, this protocol induced a fast increase in the average E-to-I synaptic weight, as shown in Figure 2B. We note that the average E-to-I synaptic weight also increased (although at a much slower rate) in Figure 1B, but in that case, the E-to-I weights in the front of the seizure decreased, and the E-to-I weights behind the seizure increased so that the inhibitory neurons in the front of the seizure could not inhibit the excitatory neurons enough to stop the propagation of the seizure (see Figure 1C). In contrast, the applied FTSTS protocol increased the strength of all the connections between the excitatory neurons and inhibitory neurons, which is shown in Figure 2E in the form of a heat map. This increase in the synaptic strengths erased the pattern of the synaptic connections induced by the seizure and thus reduced the predisposition of the neocortical network to spontaneous seizures, as shown in Figure 2C. This highlights the ability of the inverted-standard FTSTS protocol in harnessing the E-to-I synaptic plasticity of the neocortical network to suppress the prevalence of spontaneous seizures.

### Exact Reverse of the Inverted-Standard FTSTS Enhances Seizure Prevalence

We applied the exact reverse of the inverted-standard FTSTS protocol at the 10 sec mark for 5 sec after initiating the seizure in the model, which we call standard FTSTS (see Figure 3D), to the *in silico* seizure model to investigate whether our protocol could enhance the prevalence of spontaneous seizures (see Section 4 for the details of the standard FTSTS protocol and Table 1 for the standard FTSTS parameters). Compared to the inverted-standard FTSTS protocol, our standard FTSTS protocol consists of the excitatory population pulses having the polarity of 1 (i.e., the negative phase of the biphasic pulse followed by the positive phase) and the inhibitory population pulses having the polarity of −1 (i.e., the positive phase of the biphasic pulse followed by the negative phase). In this setting, the applied FTSTS protocol forces more neurons to fire in a temporal pattern of post-synaptic (inhibitory) neurons before pre-synaptic (excitatory) neurons. This leads to decrease in the average E-to-I synaptic weight of the network (see Figure 3B), lowering the activation of the inhibitory neurons during the seizure and increases the chances of the spontaneous emergence of future episodes of seizure (Figure 3C).

**Figure 3:**
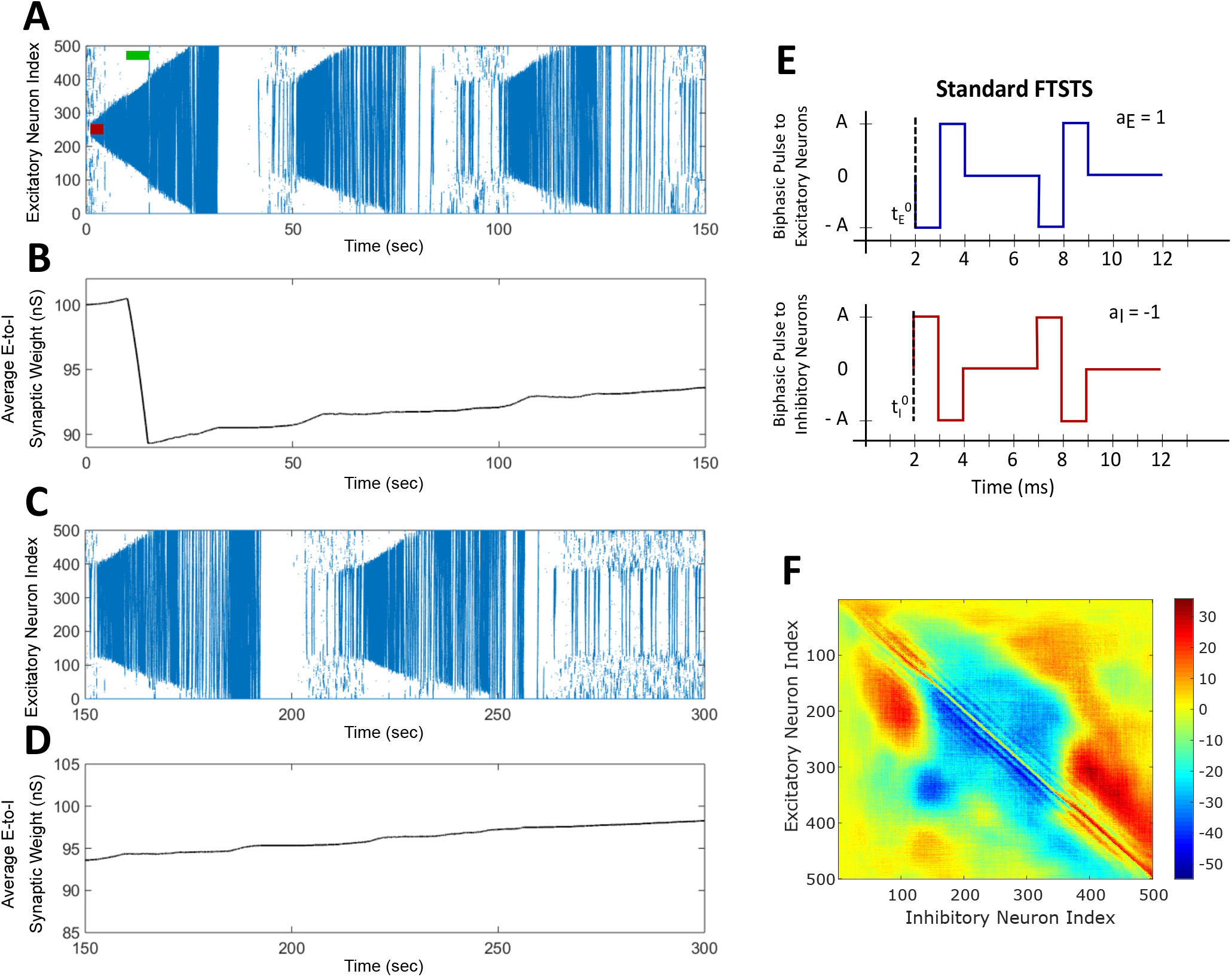
The Standard FTSTS protocol enhances the prevalence of spontaneous seizures. A seizure initiating input applied for 3 sec (red-bar) to *in silico* seizure model initiated a seizure that begins to propagate through the network. The standard FTSTS protocol was applied at 10 sec. (A) shows that the standard FTSTS protocol further enhanced the prevalence of spontaneous seizures. (B) The standard FTSTS protocol decreased the average synaptic **(B)** while it was applied. **(C)** shows the appearance of more episodes of seizure after the initial application of the standard FTSTS protocol. **(D)** shows the change in the average excitatory-to-inhibitory synaptic weight. **(E)** shows the standard FTSTS stimulation protocol where the polarity of the biphasic pulse delivered to the excitatory population is *a*_*E*_ = −1 and the polarity of the biphasic pulse delivered to the inhibitory population is *a*_*I*_ = 1. **(F)** shows the percent change in the excitatory-to-inhibitory synaptic weights at the end of the simulation (*t* = 150 sec in **(A)**).

In fact, after the FTSTS protocol, the duration of the initial seizure was much longer and the second seizure was much larger compared to the two small seizures observed in Figure 1A. The final change in the average E-to-I synaptic weight at the end of the simulation is shown in Figure 3F in the form of a heat map. Again, the synapses projecting from E-to-I neurons in front of the seizure were weakened, while synapses projecting behind the seizure were strengthened (similar to the case where the standard FTSTS protocol was not applied, see Figure 1C). This change in the neural synaptic weight structure made the neocortical network prone to spontaneous seizure in the future.

Clearly, the proposed inverted-standard and standard FTSTS protocols can control the epileptic seizures effectively by modulating the average E-to-I synaptic weight of the neocortical network. Next, we systematically investigated the effect of the FTSTS parameters on ints efficacy in suppressing/enhancing seizures.

### Effect of the Pulse Amplitude on the FTSTS efficacy

We considered four FTSTS protocols (see Table 1), which we describe in detail in the Materials and Methods Section 4. Briefly, we define our FTSTS pulse based on its polarity. We say that the FTSTS pulse has a polarity of +1 if the biphasic pulse begins with the negative amplitude. Similarly, if the biphasic pulse begins with the positive amplitude, we say that it has a polarity of −1. Based on this definition, we defined four FTSTS protocols: (1) the standard FTSTS in which the neurons in the excitatory population receive a biphasic stimulation pulse with polarity of +1 and the neurons in the inhibitory population receive a biphasic stimulation pulse with polarity of −1; (2) the inverted-standard FTSTS protocol in which the neurons in the excitatory population receive a biphasic stimulation pulse with polarity of −1 and the neurons in the inhibitory population receive a biphasic stimulation pulse with polarity of +1; (3) the mirrored FTSTS protocol in which the neurons in both the excitatory and inhibitory populations receive a biphasic stimulation pulse with polarity of +1; and (4) the inverted-mirrored FTSTS protocol in which the neurons in both the excitatory and inhibitory populations receive a biphasic stimulation pulse with polarity of −1.

We initiated a seizure in the neocortical model using the protocol described in the previous section (i.e., 200 pA applied at 1 sec mark for 3 sec to 50 excitatory neurons). We applied our four FTSTS protocols at the 10 sec mark for a duration 5 sec, where we kept the stimulation parameters same as those given in Table 1 except the amplitude of the pulses. We varied the amplitude of the FTSTS protocols between 1 nA and 2.5 nA. The efficacy of the applied FTSTS protocol was determined in terms of the rate of change of the average E-to-I synaptic weight in the neocortical-seizure model, which we defined as the mean of the change in the average E-to-I synaptic weight over the 5 sec window of the FTSTS application. Figure 4 shows the rate of change in the average E-to-I synaptic weight of the neocortical network for each corresponding FTSTS protocol amplitude for the inverted-standard, standard, mirrored, and inverted-mirrored.

**Figure 4:**
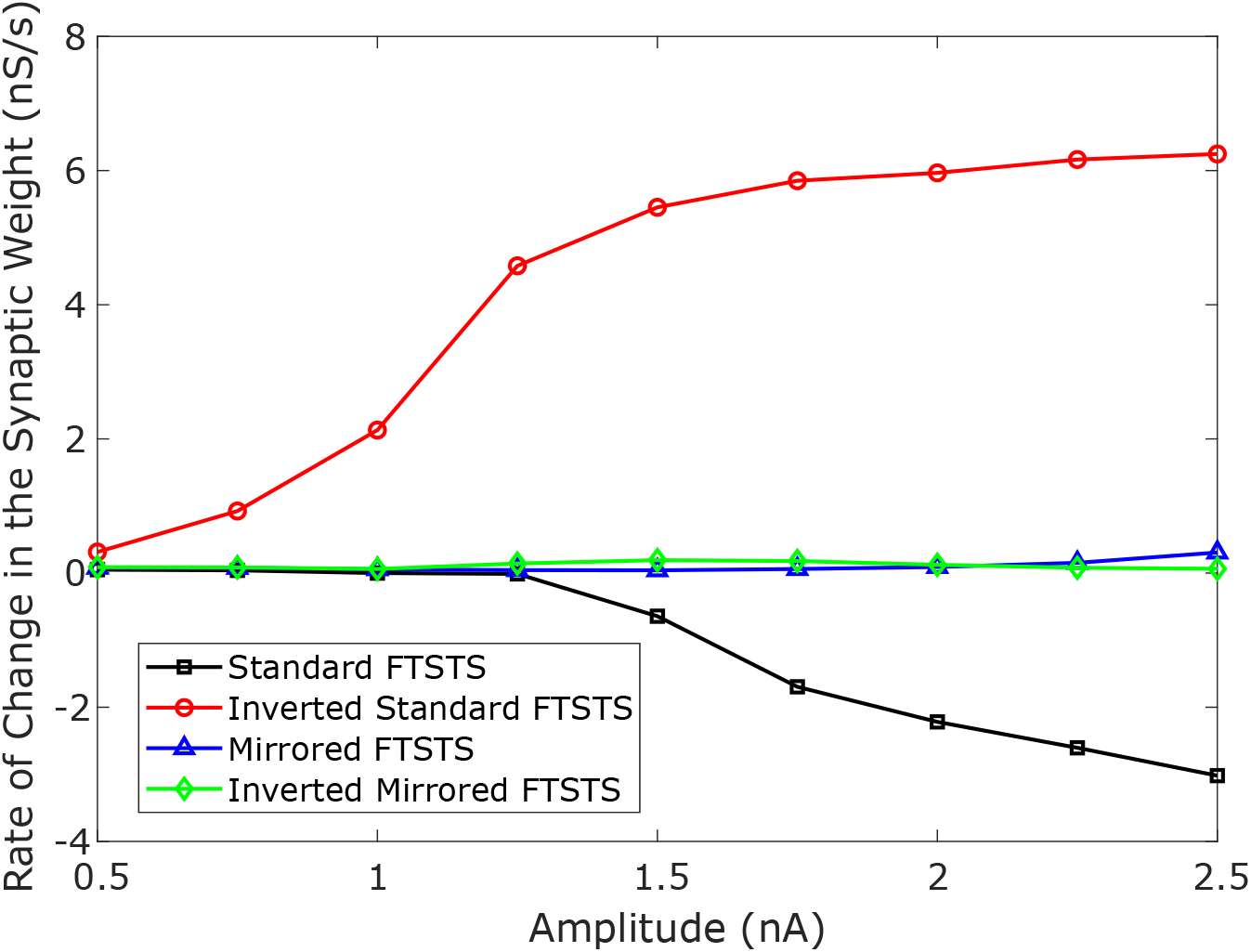
Effect of the pulse amplitude on the efficacy of FTSTS protocols. The amplitude of the FTSTS protocol was varied from 1 *nA* to 2.5 *nA*. We measured the induced change in the average synaptic weight at each pulse amplitude with a standard FTSTS polarity where *a*_*E*_ = 1 & *a*_*I*_ = −1 (black-square), inverted-standard FTSTS polarity where *a*_*E*_ = −1 & *a*_*I*_ = 1 (red-circle), mirrored FTSTS polarity where *a*_*E*_ = 1 & *a*_*I*_ = 1 (blue-triangle), and inverted-mirrored FTSTS polarity where *a*_*E*_ = −1 & *a*_*I*_ = −1 (greed-diamond).

As shown in Figure 4, the standard FTSTS protocol led to a negative rate of change in the average E-to-I synaptic strength for all amplitudes greater than 1.25 nA. As we increased the pulse amplitude, more neurons fired their action potentials in a temporal pattern of post-before-pre which led to a larger negative rate of change in the average E-to-I synaptic strength. Since the negative rate of change in the average E-to-I synaptic strength correlates positively with the increased prevalence of seizures, our simulation result suggests that applying the standard FTSTS protocol at large pulse amplitude will effectively increase the prevalence of epileptic seizure (see Figure 5).

**Figure 5:**
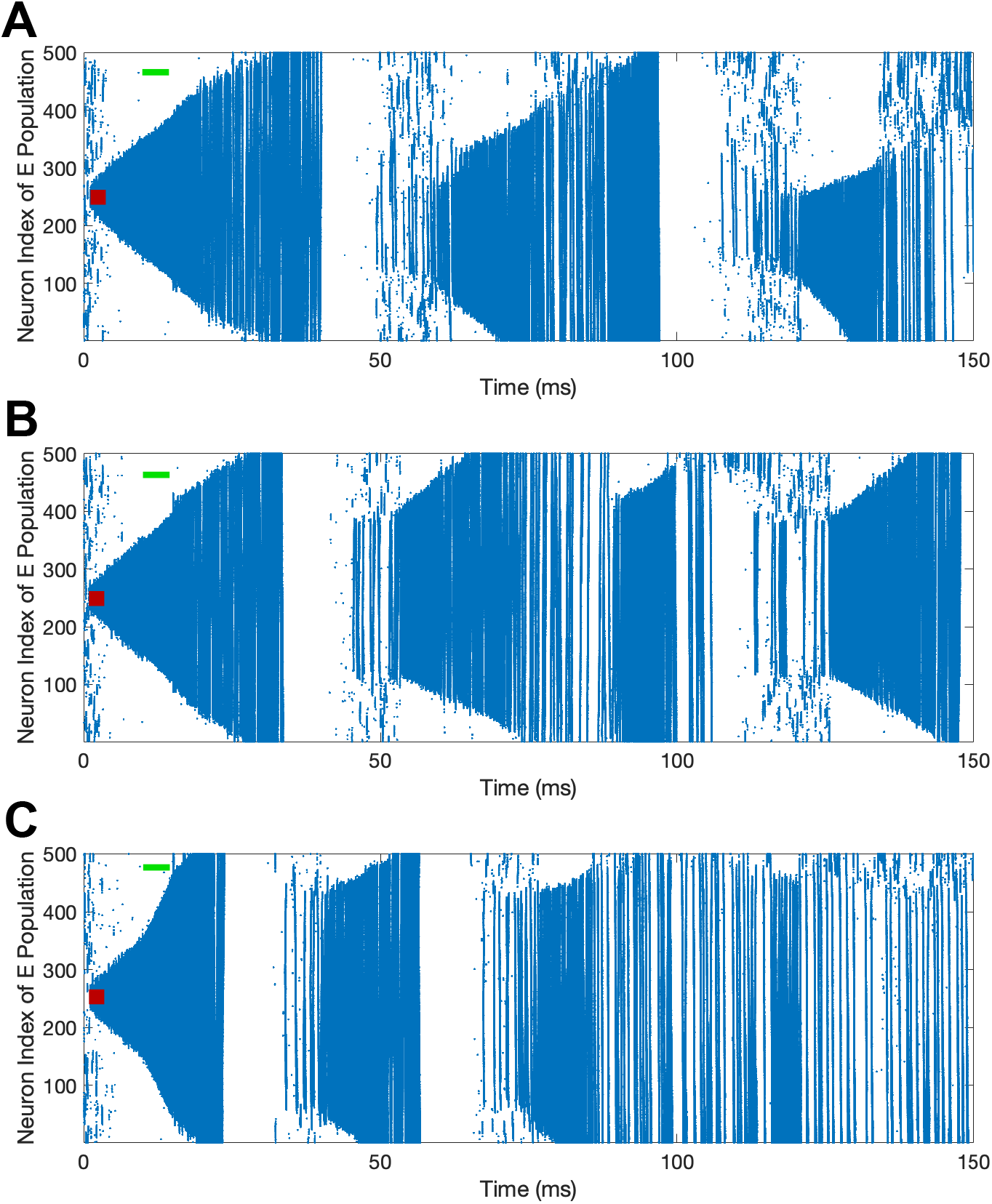
The increase in the pulse amplitude of the standard FTSTS protocol enhances the prevalence of spontaneous seizures in the *in silico* seizure model. The standard FTSTS protocol was applied for 5 sec at *t* = 10 sec with an amplitude of 1.5 nA **(A)**, 2 nA **(B)**, and 3 nA **(C)**.

The inverted-standard FTSTS protocol led to an increasing positive rate of change in the average E-to-I synaptic strength with the amplitude (see Figure 4). This increase in the average E-to-I synaptic strength was saturated for amplitude greater than 2 nA. Since the positive rate of change in the average E-to-I synaptic strength positively correlates to the suppression of seizures, our simulation result suggests that applying the inverted-standard FTSTS protocol at large pulse amplitude will effectively decrease the prevalence of epileptic seizure (see Figure 6).

**Figure 6:**
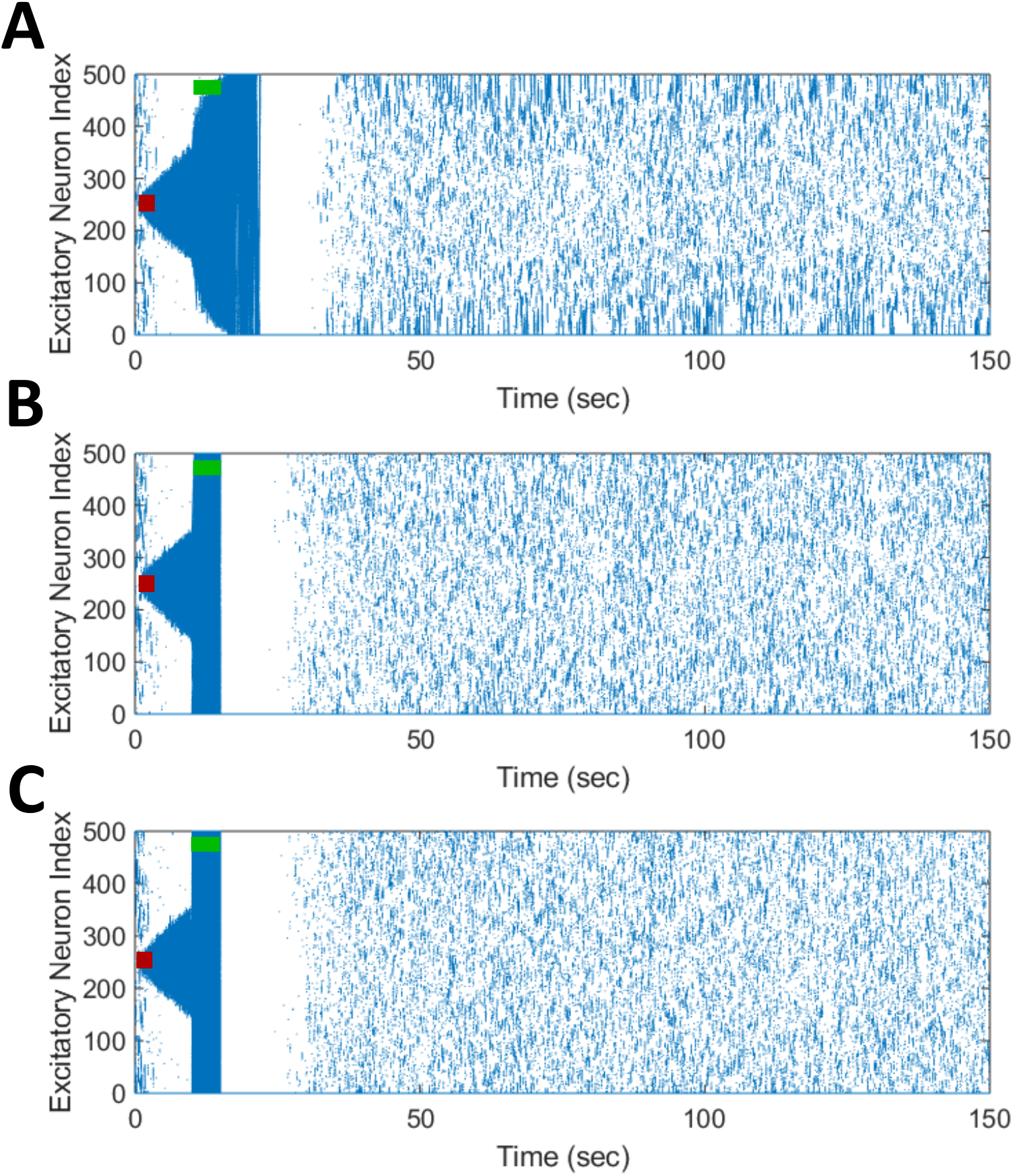
The increase in the pulse amplitude of the inverted-standard FTSTS protocol suppresses the prevalence of spontaneous seizures in the *in silico* seizure model. The inverted-standard FTSTS protocol was applied for 5 sec at *t* = 10 sec with an amplitude of 1 nA **(A)**, 1.25 nA **(B)**, and 2 nA **(C)**.

Since the same pulse was applied to the excitatory and inhibitory populations in mirrored FTSTS and mirrored-inverted FTSTS protocols (see Table 1), there was no temporal difference in the spike times induced by the FTSTS protocols. As a result, as shown in Figure 4, both the mirrored FTSTS protocol and inverted-mirrored FTSTS protocol failed to induce any changes in the average E-to-I synaptic weight of the neocortical network (shown in terms of the rate of change in the average E-to-I synaptic weight).

In conclusion, our results show that the standard FTSTS protocol and the inverted-standard FTSTS protocol at a higher amplitude and in the absence of the train-offset time are most effective in increasing and suppressing the prevalence of epileptic seizures, respectively, in this *in silico* seizure model.

### Simultaneous Effect of the Train-Offset Time and Amplitude on the FTSTS Efficacy

We varied the train-offset time of the FTSTS protocol (defined as the time difference between the start time of the inhibitory population pulse and the start time of the excitatory population pulse, also see Figure 14 in Section 4) at various pulse amplitudes for each of the four FTSTS protocols (i.e., inverted-standard, standard, mirrored, and inverted-mirrored). Similar to previous sections, we induced a seizure in the model by applying the seizure initiating input current (200 pA applied at 1 sec mark for 3 sec to 50 excitatory neurons) and applied the FTSTS protocols to the model at the 10 s mark for a duration of 5 sec. For all four FTSTS protocol, we kept the stimulation parameters same as given in Table 1 except we varied the pulse amplitude between 1 nA and 2.5 nA and the train-offset time between 0 ms and 12 ms. We noted that the train-offset time completes a cycle every 12 ms for the specified pulse width and pulse interval. Thus, a train-offset time of 0 ms is equal to a train-offset time of 12 ms and a train-offset time of −1 ms is equal to a train-offset time of 11 ms. This allowed us to construct a surface plot of the full train-offset cycle vs pulse amplitude for each pulse-pair polarity.

Figure 7A shows a surface plot displaying the effect of the pulse amplitude versus train-offset time on the rate of change in the average E-to-I synaptic weight (i.e., the efficacy of the protocol). As shown in this figure, the standard FTSTS protocol with a train-offset time of 0.5 ms or −11.5 ms led to the maximum decrease in the average E-to-I synaptic weight over the span of the considered amplitude parameters with a higher amplitude producing a larger decrease. A train-offset time of 0.5 ms (or −11.5 ms) for the standard FTSTS polarity occurred when the positive portion of the inhibitory population pulse began 0.5 ms before the positive portion of the excitatory population pulse. Based on the above result of the optimal train-offset time to decrease the average E-to-I synaptic weight, one would have expected the optimal train-offset for increasing the average E-to-I synaptic weight to be the reverse order. This would correspond to the positive portion of the excitatory population pulse arriving 0.5 ms before the positive portion of the inhibitory population pulse. Due to the asymmetrical nature of the standard FTSTS protocol (see Figure 3E), this would correspond to the train-offset time of 1.5 ms (−10.5 ms).

**Figure 7:**
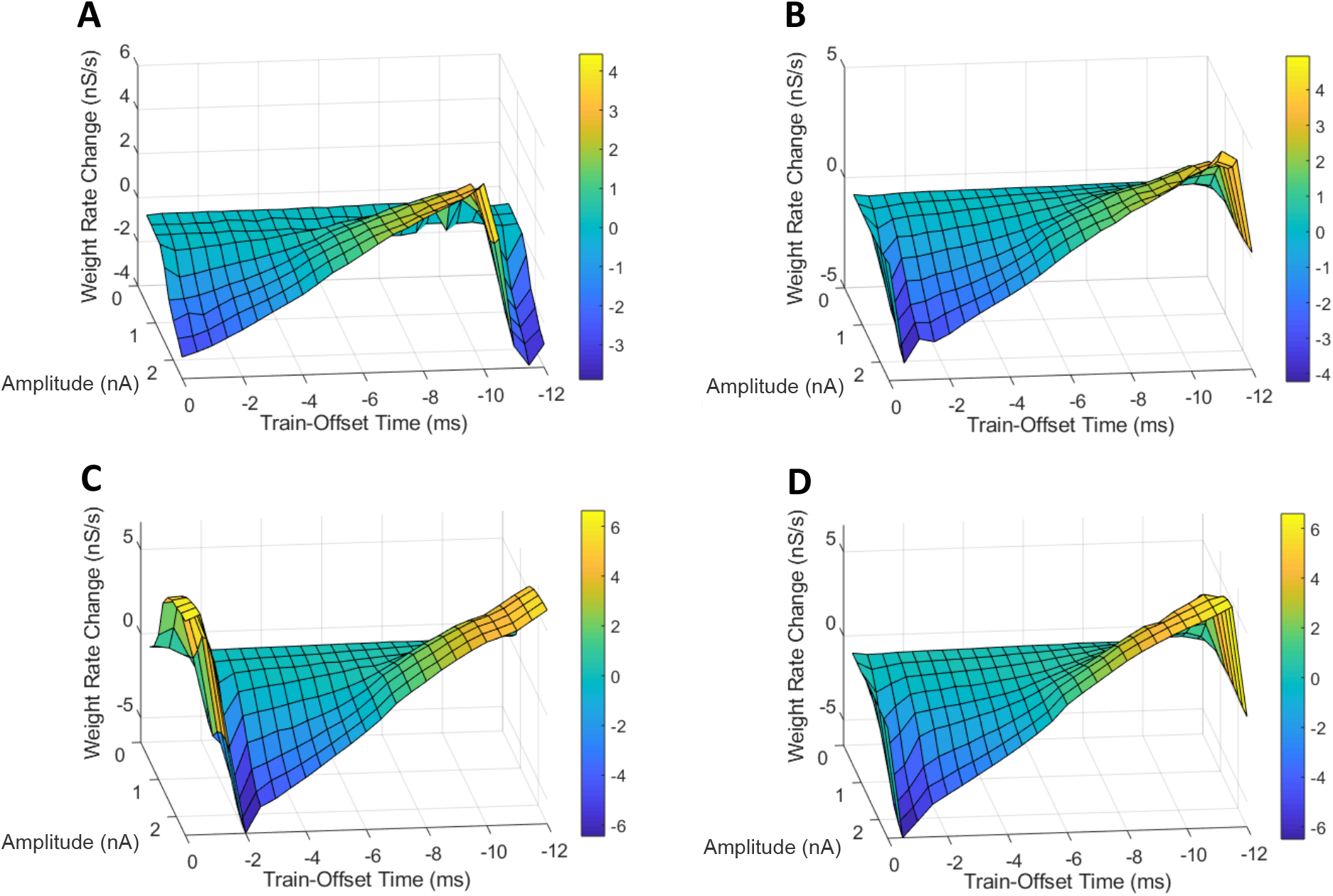
Effects of the train-offset time and amplitude on the efficacy of the FTSTS protocols in suppressing or enhancing the prevalence of epileptic seizure. The efficacy of the FTSTS was measured for various amplitudes and phase differences. The change in the average synaptic weight from the 5 sec application of the FTSTS protocol during a seizure. The effect of the amplitude and phase difference of the FTSTS protocol for the four possible polarities **(A)** the standard FTSTS polarity (*a*_*E*_ = 1 and *a*_*I*_ = −1), **(B)** mirrored FTSTS polarity (*a*_*E*_ = 1 and *a*_*I*_ = 1), **(C)** inverted standard FTSTS polarity (*a*_*E*_ = −1 and *a*_*I*_ = 1), and **(D)** inverted mirrored FTSTS polarity (*a*_*E*_ = −1 and *a*_*I*_ = −1) on FTSTS efficacy were considered.

However, we did not observe this optimal train-offset time in our simulations. We observed that the optimal train-offset time to increase the average E-to-I synaptic weight was 2 ms (−10 ms) with a larger amplitude producing greater increase. A possible reason for this deviation was that we defined the positive polarity (+1) of the biphasic pulse as the stimulating pulse starting with the negative portion of the bi-phasic pulse, which is then followed by the positive portion. Therefore, the positive portion of the excitatory population pulse must overcome the inhibition of the excitatory neuron population by the negative portion of the pulse, which resulted in a slower response of the excitatory neurons. This required the inhibitory neurons to be stimulated a little later to achieve a more favorable temporal spike-time pattern between the excitatory and inhibitory populations. Since the inhibitory population pulse had a negative polarity (-1), the positive portion of the pulse led the negative portion. Therefore, the positive portion of the bi-phasic pulse did not have to overcome any inhibition of the neuron population by its negative counterpart. While our investigation of the amplitude and train-offset time parameter space highlighted the optimal amplitude and train-offset parameters to decrease or increase the average E-to-I synaptic weight of the network, it also highlighted the importance of the FTSTS pulse polarity.

Figure 7C shows a surface plot displaying the effect of the pulse pair train-offset time and pulse amplitude of the inverted-standard FTSTS protocol on the rate of change in the average E-to-I synaptic weight of the neocortical network. For all train-offset time, the magnitude of the efficacy of the inverted-standard FTSTS protocol increased with the pulse amplitude. As shown in Figure 7C, the most effective train offset time to increase the average E-to-I synaptic weight was −0.5 ms (11.5 ms) and to decrease the average E-to-I synaptic weight was −2 ms (10 ms). Similar to the standard FTSTS protocol, we observed asymmetry in the optimal train-offset time for the inverted-standard FTSTS protocol. Since the pulses delivered to the inhibitory population led with the negative portion of the biphasic pulse, the positive portion of the biphasic pulse must overcome this initial inhibition caused by the negative portion of the pulse. Since the inhibitory population is required to fire before the excitatory population, the train-offset time must increase to accommodate for the delay in the spike times.

Figures 7B and 7D show the surface plots displaying the efficacy of the mirrored and inverted-mirrored FTSTS protocols, respectively, at various train-offset time and pulse amplitude. As noted in these figures, the magnitude of the efficacy of the FTSTS protocol increased with the pulse amplitude. We found that the most effective train-offset time to increase the average E-to-I synaptic weight was −1 ms and −0.5 ms (11 ms and 11.5 ms) for mirrored and inverted mirrored, respectively. While the most effective train-offset time was −11.5 ms (0.5 ms) to decrease the average E-to-I synaptic weight for both protocols. Unlike the standard FTSTS protocol and the inverted-standard FTSTS protocol, we found that the optimal train-offset time was almost symmetric in these cases. Since the mirrored FTSTS protocol ends with the excitatory portion of the biphasic pulse, the excitatory neurons continue to fire during the interpulse interval. This shifts the optimal train-offset time slightly to −1 ms to increase the synaptic weight and reduced the efficacy. In the case of the inverted FTSTS protocol, it ends with the inhibitory portion of the FTSTS protocol, which stops the neuronal firing. This creates the symmetric train offset times of −0.5 ms and 0.5 ms to increase and decrease the average synaptic weight respectively. We further observed that the inverted-mirrored FTSTS protocol was more effective in modulating the average E-to-I synaptic weight compared to the mirrored FTSTS protocol.

In conclusion, our simulation results highlight the train-offset time as a critical FTSTS parameter to be optimized to achieve optimal efficacy of the applied FTSTS protocol in decreasing/increasing the prevalence of epileptic seizures in an *in silico* seizure model.

### Simultaneous Effect of the Pulse Frequency and the Train-Offset Time on the FTSTS Efficacy

In this section, we investigated how the FTSTS pulse frequency and the pulse pair train-offset time of the delivered FTSTS pulses simultaneously affect the efficacy of the four FTSTS protocols in modulating the average E-to-I synaptic weight of the neocortical network in the *in silico* seizure model. Similar to previous sections, we induced a seizure in the model by applying the seizure initiating input current (200 pA applied at 1 sec mark for 3 sec to 50 excitatory neurons) and applied the FTSTS protocols to the model at the 10 sec mark for a duration of 5 sec. For all four FTSTS protocol, we kept the stimulation parameters the same as those given in Table 1 except we varied the pulse frequency and the train-offset time. Specifically, we considered seven different stimulation frequency (143 Hz, 83 Hz, 45 Hz, 31 Hz, 24 Hz, 19 Hz, and 9.8 Hz) and varied the train-offset time between −3.5 ms and 3.5 ms. Figure 8A shows our simulation results in the form of a surface plot displaying the efficacy of the standard FTSTS protocol, measured in the form of the rate of change in the average E-to-I synaptic weight, at various stimulation frequency and the pulse pair train-offset time. As shown in this figure, the efficacy of the standard FTSTS protocol enhanced with the increase in the stimulation frequency for all train-offset time with an exception at 143 Hz. Particularly, when we increased the stimulation frequency from 83 Hz to 143 Hz in our simulation, the efficacy of the standard FTSTS protocol decreased at high values of the train-offset time. A potential explanation for this observation is as follows. The highest frequency that we considered was 143 Hz, which corresponds to a pulse interval of 5 ms. This short inter-pulse interval not only forced the presynaptic (excitatory) neurons before the postsynaptic (inhibitory) neurons but also the postsynaptic (inhibitory) neurons before the presynaptic (excitatory) neurons with spike time differences that were within the timescale of the Hebbian STDP. For example, consider the case of the standard FTSTS protocol with a train-offset time of 0 ms and a pulse interval of 5 ms (i.e., 143 Hz) in Figure 8A. This protocol not only induces a pre-post firing with a pre-post spike time difference of 1 ms in the neocortical network but also induces a post-pre firing with a post-pre spike time difference of 5 ms. This closeness of the spike time difference in both pre-post and post-pre firings counteracted the change mediated by the Hebbian STDP (timescale of 15 ms) in the E-to-I synaptic weights and reduced the protocol’s efficacy in increasing or decreasing the average E-to-I synaptic weight of the neocortical network. This highlights that both the lower and higher frequencies are less efficient.

**Figure 8:**
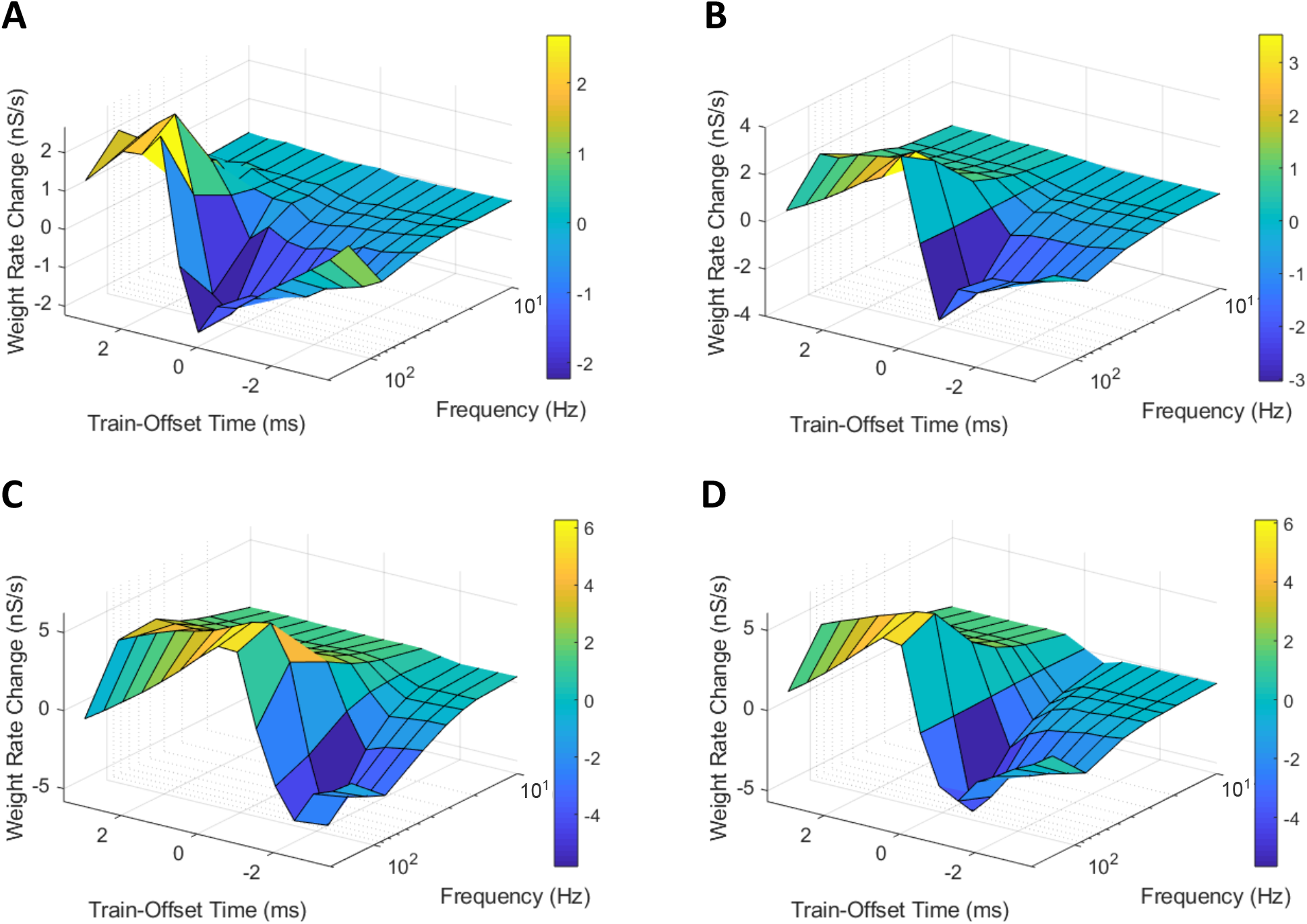
Effects of the pulse frequency and train-offset time on the FTSTS protocols in suppressing or enhancing the prevalence of epileptic seizure. We measured the efficacy of the FTSTS protocol at different frequencies, train-offset times, and polarity pairs. The efficacy was measured as the rate of change in the average synaptic weight over the 5 second interval the FTSTS was applied for the **(A)** standard (*a*_*E*_ = −1 and *a*_*I*_ = − 1), **(B)** mirrored (*a*_*E*_ = −1 and *a*_*I*_ = 1), **(C)** inverted standard (*a*_*E*_ = −1 and *a*_*I*_ = 1), and **(D)** inverted mirrored (*a*_*E*_ = −1 and *a*_*I*_ = − 1) FTSTS protocols.

For the standard FTSTS protocol, we found that the optimal frequency was 83 Hz, which balanced the increased efficacy from more desired forced spike time difference (post-pre firing) and decreased efficacy from the undesired forced spike time difference (pre-post firing) at higher frequencies. Particularly, we found that the optimal stimulation frequency and train-offset time for the standard FTSTS polarity to increase the average E-to-I synaptic weight was 83 Hz and 2 *ms*, respectively. Additionally, the optimal stimulation frequency and train-offset time to decrease the average E-to-I synaptic weight was 143 Hz and 0.5 *ms*, respectively. Although the inherent seizure frequency in the *in silico* seizure model was in the Theta range (5-7 Hz), we found that the optimal FTSTS frequency was independent of the inherent seizure frequency and suppressed the LFP power in all frequency ranges (not shown here).

Next, we examined the effect of the frequency and the train-offset time on the efficacy of the inverted-standard FTSTS protocol. Figure 8C shows the average E-to-I synaptic weight change at different frequency and train-offset times for the inverted-standard FTSTS polarity. The efficacy of the FTSTS protocol increased with the frequency of stimulation except at the highest two frequencies. At these frequencies, the efficacy decreased for all train-offset values when the frequency was increase from 83 *Hz* to 143 *Hz*. Again, this is due to the undesired forced spiketimes counteracting the desired forced spike-time difference at the shorter pulse interval (i.e, higher frequency). The optimal stimulation frequency and train-offset time for the inverted-standard FTSTS polarity to increase the average E-to-I synaptic weight was 83 Hz and −0.5 *ms*, respectively. Furthermore, the optimal stimulation frequency and train-offset time to decrease the average E-to-I synaptic weight was 83 Hz and −2 *ms*, respectively.

Then, we analyzed the effect of frequency and train-offset time on the efficacy of the mirrored FTSTS protocol. Figure 8B shows that the efficacy of the FTSTS protocol increases with the frequency except for the two highest frequencies. We observed a decrease in the average E-to-I synaptic weight rate change when the frequency was increased from 83 Hz to 143 Hz for all train-offset times except −0.5 *ms* and 0.5 *ms*. This corresponds to the positive portion of the inhibitory biphasic pulse arriving half of a millisecond before and after the positive portion of the excitatory biphasic pulse. We observed the same trend for the standard FTSTS polarity. The general decrease in the efficacy of the FTSTS protocol going from a stimulation frequency of 83 Hz to 143 Hz was from the undesired forced spike-time difference between the two neuron populations counteracting the desired forced spike-time difference at the higher frequency. Two exceptions to this trend occurred when the difference between desired and undesired forced spike-times of the two neuron populations was the greatest (−0.5 *ms* and 0.5 *ms*). The optimal stimulation frequency and train-offset time for the mirrored FTSTS polarity to increase the average E-to-I synaptic weight was 143 Hz and 0.5 *ms*, respectively. Additionally, the optimal stimulation frequency and train-offset time to decrease the average E-to-I synaptic weight was 83 Hz and −0.5 *ms*, respectively.

Finally, we considered the effect of frequency and train-offset time on the efficacy of the inverted-mirrored FTSTS protocol. This is shown in Figure 8D. The magnitude of the average E-to-I synaptic weight rate change (efficacy) increased with the stimulation frequency up to a frequency of 83 *Hz*. We observed when the frequency was increased from 83 *Hz* to 142 *Hz* the efficacy decreased for all train-offset times. Similar to the other three polarities, this occurred due to the induced undesired forced spike-times counteracting the desired forced spiketime difference at the shorter pulse interval (higher frequency). For the inverted-mirrored FTSTS protocol, the optimal pulse interval and train-offset time to increase the average synaptic weight was 10 *ms* (83 *Hz*) and 0.5 *ms*, respectively. Additionally, the optimal pulse interval and train-offset time to decrease the average synaptic weight was 10 *ms* (83 *Hz*) and −0.5 *ms*, respectively.

In conclusion, our simulation results show that an increase in the stimulation frequency increased the efficacy except at the highest frequency. Additionally, at the highest frequency there were less deviations from this trend with the inverted polarities. Clearly, this established the optimal polarity of the excitatory population biphasic pulse as *a*_*E*_ = −1. A possible reason for this observation is the negative polarity begins with the positive portion of the biphasic pulse. Therefore, the pulse can more quickly force the neurons in the excitatory population to fire, since it is not required to overcome any inhibition from the negative portion of the biphasic pulse. These results highlight the importance of the ability of FTSTS to force the neurons to fire during the positive portion of the bi-phasic pulse. Therefore, optimizing the remaining FTSTS parameter, pulse width, may further increase the efficacy of the protocol.

### Simultaneous Effect of the Pulse Width and the Train-Offset Time on the FTSTS Efficacy

In this section, we present our results on the simultaneous effect of the pulse width and the pulse pair train-offset time on the efficacy of all four FTSTS protocols in suppressing or enhancing the seizure prevalence. Similar to previous sections, we induced a seizure in the model by applying the seizure initiating input current (200 pA applied at 1 sec mark for 3 sec to 50 excitatory neurons) and applied the FTSTS protocols to the model at the 10 sec mark for a duration of 5 sec. For our investigation, we considered the pulse width values between 1 ms and 5 ms and the train-offset time between −10 ms to 10 ms. All other stimulation parameters were same as given in Table 1.

We first considered the standard FTSTS protocol and investigated the effect of the pulse width at different train-offset time on the efficacy of this protocol in modulating the average E-to-I synaptic weight of the neocortical network. Figure 9A shows the simultaneous effect of the pulse width and the train-offset time on the efficacy of the standard FTSTS protocol in decreasing the average E-to-I synaptic weight. In general, the optimal train-offset time to decrease the average E-to-I synaptic weight occurred at *W* + 1 *ms*, where *W* is the pulse width. This timing attempted to force the inhibitory neurons to fire 1 *ms* before the excitatory neurons by delivering the positive portion of the bi-phasic pulse to the inhibitory population 1 *ms* before the positive portion of the pulse delivered to the excitatory population. The exception to this general train-offset timing pattern occurred at pulse widths greater than 3 *ms*. For pulse widths of 4 *ms* and 5 *ms*, the optimal train-offset time occurred at 8 *ms*.

**Figure 9:**
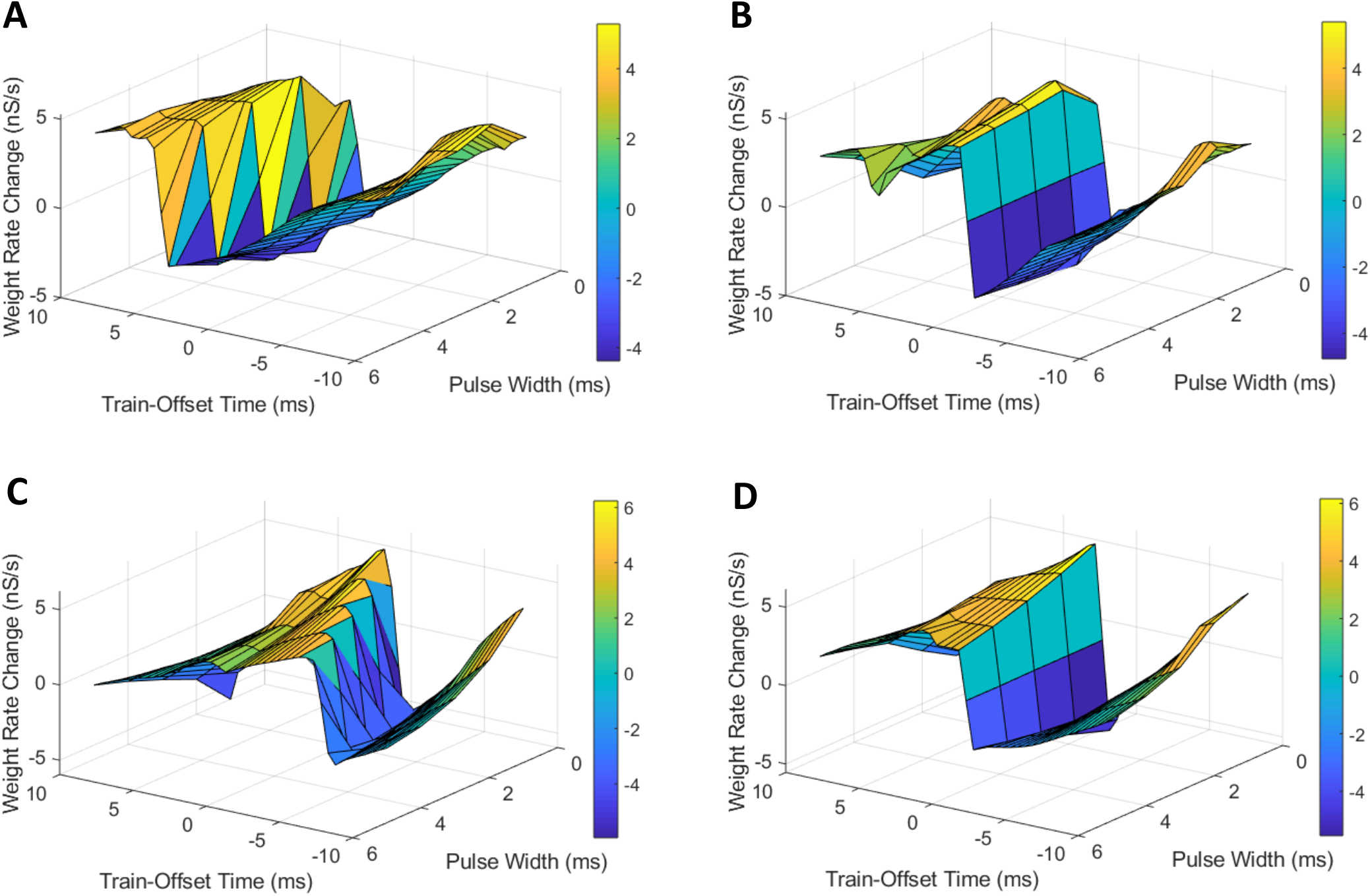
Effects of the pulse width and train-offset time on the FTSTS protocols in suppressing or enhancing the prevalence of epileptic seizure. We measured the efficacy of the FTSTS protocol at different pulse widths, train-offset times, and polarity pairs. The efficacy was measured as the rate of change in the average synaptic weight over the 5 second interval the FTSTS was applied for the **(A)** standard (*a*_*E*_ = −1 and *a*_*I*_ = − 1), **(B)** mirrored (*a*_*E*_ = −1 and *a*_*I*_ = 1), **(C)** inverted-standard (*a*_*E*_ = −1 and *a*_*I*_ = 1), and **(D)** inverted-mirrored (*a*_*E*_ = −1 and *a*_*I*_ = − 1) FTSTS protocols.

Then, we examined the efficacy of the parameters pulse width and train-offset of the inverted-standard FTSTS polarity. The optimal train-offset time at each pulse width to increase or decrease the average E-to-I synaptic weight is shown in Figure 9C as the peak or trough, respectively. The efficiency of the inverted-standard FTSTS protocol decreased with the pulse width for both increasing and decreasing the average E-to-I synaptic weight. The trend of the optimal train-offset time for each pulse width to increase the average E-to-I synaptic weight is −*W* + 0.5 *ms*, where *W* is the pulse width. This resulted in the positive portion of the bi-phasic pulses delivered to the excitatory population arriving 0.5 *ms* before the positive portion delivered to the inhibitory population. Due to the opposite polarities of the pulses delivered to each population, the negative portion of the bi-phasic pulse arrived before the positive portion of the pulse. This prevented the inhibitory neurons from firing before the excitatory neurons. Additionally, the negative portion of the bi-phasic pulse delivered to the excitatory population arrived mostly after positive portion was delivered to the inhibitory population, which prevented the excitatory neurons from firing after the inhibitory neurons. The optimal train-offset time to decrease the average E-to-I synaptic weight followed a general trend of −*W* − 1 *ms* for a pulse width less than 4 *ms* and a trend of *W* − 1.5 *ms* for pulse widths greater than or equal to 4 *ms*. For pulse widths less than 4 *ms*, the positive portion of the bi-phasic pulse delivered to the inhibitory population arrived 1 *ms* prior to the positive portion delivered to the excitatory population. As the pulse width increased, the optimal train-offset time decreased to −*W* − 1.5 *ms*. This may result from the increased inhibition of the inhibitory neurons by the negative portion of the bi-phasic pulse as the pulse width increased, which the positive portion of the bi-phasic must overcome to force the inhibitory neurons to fire. Therefore, the inhibitory neurons responded slower and required a larger gap between when the excitatory and inhibitory neurons are stimulated to ensure that most of the inhibitory neurons fire before the excitatory neurons.

Next, we determined the efficacy of the pulse width and train-offset time on the efficacy of the mirrored FTSTS polarity. The symmetric polarities result in optimal train-offset times to increase or decrease the average E-to-I synaptic weight that were the same across all pulse widths as shown in Figure 9B. The optimal train-offset time to increase the average E-to-I synaptic weight for all pulse widths was 1 *ms*. This resulted in the positive portion of the bi-phasic pulse delivered to the excitatory neurons arriving 1 *ms* before the positive portion delivered to the inhibitory population. Due to the symmetry of the polarity, the neurons consistently respond to the positive portion of the pulse train in both of the neuron populations. The optimal pulse width and train-offset time to increase the average E-to-I synaptic weight was 2 *ms* and 1 *ms*, respectively. Additionally, the best train-offset time for all pulse widths to decrease the average E-to-I synaptic weight was −0.5 *ms*. Therefore, the positive portion of the bi-phasic pulse delivered to the inhibitory population arrived 0.5 *ms* before the portion delivered to the excitatory neurons. Again, the symmetry of the polarities ensured the neurons in both of the populations responded on the same timescale. The optimal pulse width and train-offset time to decrease the average E-to-I synaptic weight of the network was 2 *ms* and −0.5 *ms*, respectively.

Finally, we analyzed how the pulse width and train-offset time parameters influenced the efficacy of the inverted-mirrored FTSTS protocol. The surface plot of the change in average E-to-I synaptic weight for the different parameter combinations of pulse width and train-offset time is shown in Figure 9D. The efficacy of the inverted FTSTS protocol decreased with pulse width for both increasing and decreasing the average E-to-I synaptic weight. For pulse widths between 1 *ms* and 3 *ms*, the optimal train-offset time was 0.5 *ms*. The symmetric polarities results in the positive portion of the bi-phasic pulse delivered to the excitatory population arriving 0.5 *ms* before the positive portion delivered to the inhibitory population. Also, the negative portion following the positive portion of the biphasic pulse prevented unwanted neuron firing in both populations after forcing the neurons to fire. At the longer pulse widths considered, the train -offset times for a pulse width of 4 *ms* and 5 *ms* was 3.5 *ms* and 3 *ms*, respectively. These train-offset times result in a majority of the positive portion of the bi-phasic pulse delivered to the excitatory population arriving before the positive portion delivered to the inhibitory population. This may be due to the longer pulse width forcing unwanted firing when the positive portions of the the bi-phasic pulse overlapped, which occurred less at shorter pulse widths due to the 2 *ms* refractory period. The optimal train-offset times to decrease the average E-to-I synaptic weight was −0.5 *ms* for all pulse widths. This resulted in the positive portion of the bi-phasic pulse delivered to the inhibitory population arriving 0.5 *ms* before the positive portion delivered to the inhibitory population for all pulse widths. Due to the inhibitory input to the excitatory neurons resulting from the forcing the inhibitory neurons to fire, there was less unwanted firing when the positive portions of the bi-phasic pulses delivered to each population overlapped. Additionally, the negative portion of the bi-phasic pulse prevented unwanted firing in both populations, after forcing the neurons of the opposite population to fire. The optimal train-offset time and pulse width to decrease the average E-to-I synaptic weight was −0.5 *ms* and 1 *ms*, respectively.

In conclusion, our simulation results show a linear relationship between the optimal train-offset time and the FTSTS pulse width in increasing or decreasing the average E-to-I synaptic weight in the neocortical epileptic computational model for all FTSTS protocols except the mirrored FTSTS protocol.

### Efficacy of Inverted-Standard FTSTS Protocol in Partially Inseparable E-I Population

One potential limitation of testing our designed FTSTS protocol in animal experiments is that our electrical FTSTS protocol, described in the previous sections, requires spatially separable targeted excitatory and inhibitory populations. To address this issue, in this section, we investigated whether our inverted-standard FTSTS protocol can efficiently suppress the prevalence of spontaneous seizures in the *in silico* seizure model when a fraction of the excitatory and inhibitory neurons receives both the excitatory and inhibitory FTSTS pulses. We varied the percentage of neurons in each population that receives excitatory and inhibitory FTSTS pulses from 0% to 80%. Here, 0% means that excitatory neurons only receive excitatory FTSTS pulses and inhibitory neurons only receive inhibitory FTSTS pulses. In contrast, 80% means that 80% of randomly selected excitatory neurons also receive inhibitory FTSTS pulses, and 80% of the randomly selected inhibitory neurons also receive excitatory FTSTS pulses. In each case, we induced a seizure and applied the inverted-standard FTSTS protocol using the same protocols described in previous sections. All other stimulation parameters were the same as those given in Table 1 for the inverted-standard FTSTS protocol.

Our simulation results shown in Figure 10 indicate that our inverted-standard FTSTS protocol can increase the average E-to-I synaptic weight and thus suppress the prevalence of spontaneous seizures for the stimulation overlap up of up to 60% with a decaying efficacy. For the scenario with 80% overlap, the inverted-standard FTSTS protocol failed to prevent a seizure, which can be seen in the steady increase of the average synaptic weight at 70 sec (blue line). In conclusion, our simulation results show that if there exists a moderate level of inseparability in the excitatory and inhibitory neurons in an E-I population, our inverted-standard FTSTS protocol will still be able to control the synaptic weight and prevent a seizure in the *in silico* seizure model.

**Figure 10:**
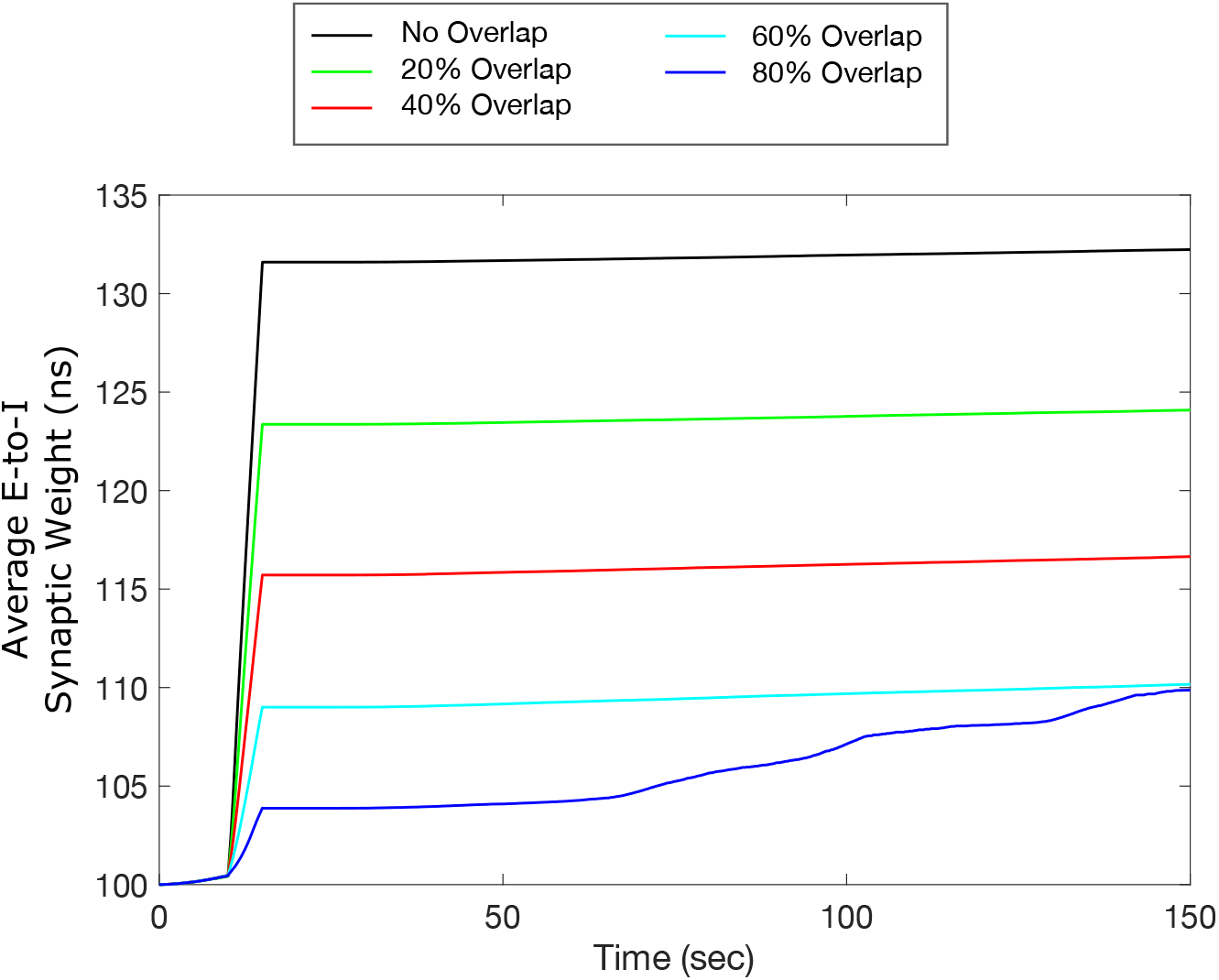
Efficacy of the inverted-standard FTSTS protocol in suppressing the prevalence of epileptic seizures where E-I population is partially separable. We applied our inverted-standard FTSTS protocol with parameters listed in Table 1 with 0% (black line), 20% (green line), 40% (red line), 60% (cyan line), and 80% (blue line) overlap of both the excitatory and inhibitory neuron population.

### Optimal FTSTS Protocol for Suppressing/Enhancing Seizure

After systematically analyzing the FTSTS pulse design parameters on the efficacy of 4 different FTSTS protocols in modulating the average E-to-I synaptic weight of the neocortical network in the previous sections, we investigated the optimal FTSTS protocol and corresponding pulse design parameters in this section. In particular, we found that the inverted-standard FTSTS protocol with biphasic pulses consisting of a pulse interval of 10 ms (or equivalently the stimulation frequency of 83 Hz), a pulse width of 1 ms, a train-offset time of −0.5 ms, and an amplitude of 2 nA is optimal to increase the average E-to-I synaptic weight of the neocortical network. It should be noted here that the rate of change in the average synaptic weight to increase the average E-to-I synaptic weight plateaued at amplitudes greater than 2 nA (see Figure 4 in Section 2). Moreover, we found that the same protocol (i.e., the inverted-standard FTSTS protocol) with biphasic pulses consisting of a pulse interval of 10 ms (or equivalently the stimulation frequency of 83 Hz), a pulse width of 1 ms, a train-offset time of −2 ms, and an amplitude greater than 2 nA is most effective in decreasing the average E-to-I synaptic weight of the neocortical-seizure model.

To demonstrate the efficacy of the optimal FTSTS protocol in suppressing and enhancing the seizure prevalence, we applied these protocols to the neocortical-seizure model at the 10 sec mark for 5 sec. Figure 11A shows our simulation results for the applied inverted-standard protocol to increase the average E-to-I synaptic weight of the neocortical network. As shown in this figure, the optimal FTSTS protocol terminated the initial seizure induced by a seizure-like input in the neocortical network immediately after the applied duration of the FTSTS protocol. Additionally, we did not observe the emergence of any spontaneous seizure over the period of 150 sec. Figure 11B shows the increase in the average E-to-I synaptic weight of the neocortical network induced by the FTSTS protocol.

**Figure 11:**
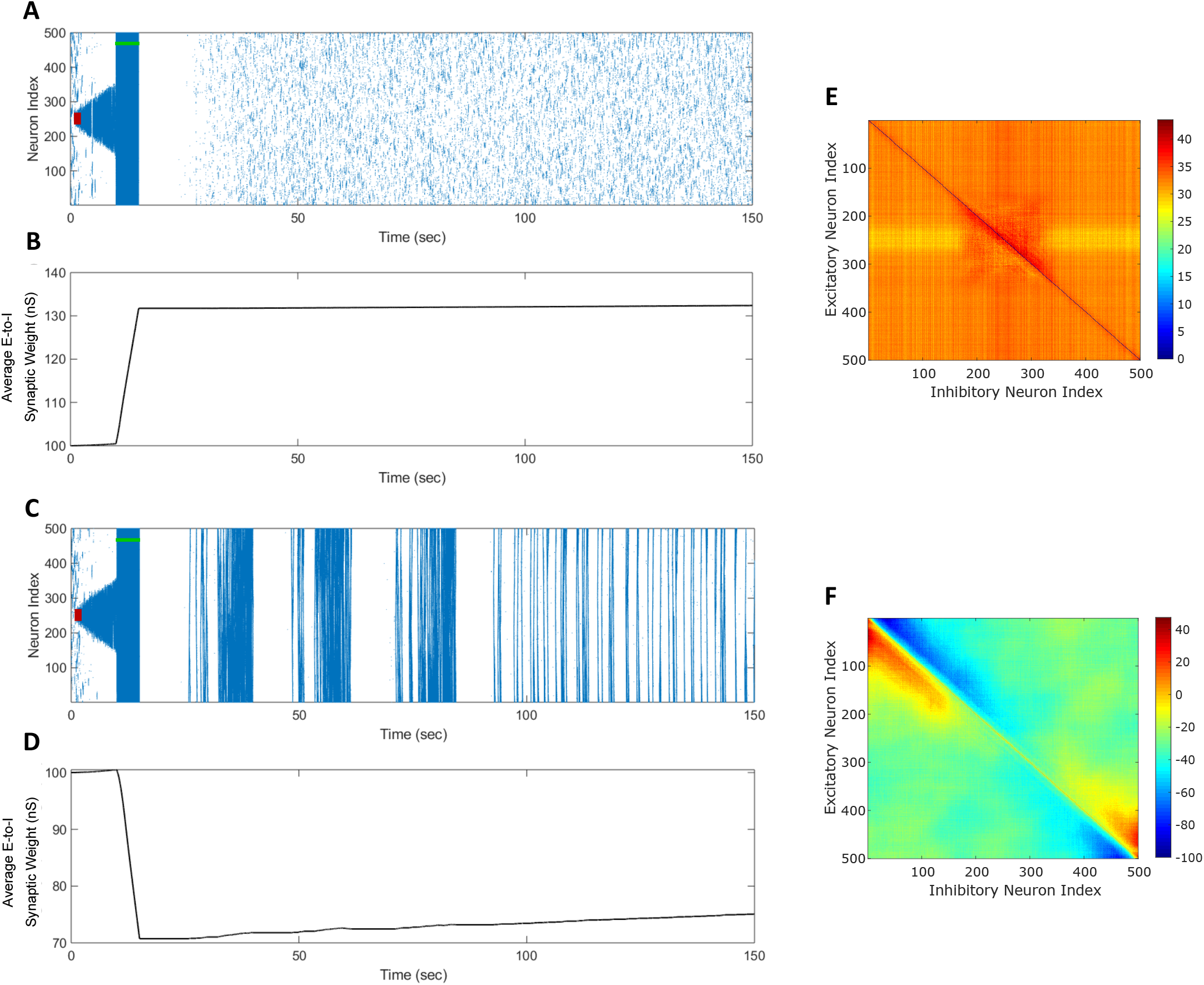
Optimal FTSTS Parameters to suppress or enhance the prevalence of the episodes of epileptic seizure. We applied our FTSTS protocol with a pulse amplitude of 2 *nA*, pulse interval of 10 *ms*, a pulse width of 1 *ms*, and an inverted standard polarity (*a*_*E*_ = −1 and *a*_*I*_ = 1). The initial seizure was induced by a seizure inducing input of 200 *pA* (red-bar). We applied a train-offset time of −0.5 *ms* to increase the average synaptic weight in **(A)** in order to stop the initial seizure and prevent future seizures. **(B)** shows the increase in the average synaptic weight induced by FTSTS protocol (green-bar). We applied a train-offset time of −2 *ms* to decrease the average synaptic weight in **(C)**, which initially stopped the first seizure but produced strong spontaneous seizure for the rest of the simulation. **(D)** shows the decrease in the average synaptic weight induced by FTSTS protocol (green-bar). **(E)** and **(F)** show the percent change in the excitatory-to-inhibitory synaptic weights at the end of the simulation in **(A)** and in **(C)**, respectively.

Figure 11C shows our simulation results for the optimal inverted-standard protocol to decrease the average E- to-I synaptic weight of the neocortical network. As shown in this figure, the optimal FTSTS protocol terminated the initial seizure induced by a seizure-like input in the neocortical network immediately after the applied duration of the FTSTS protocol. Shortly after the termination of the first episode of seizure, the changes in the E-to-I synaptic weights induced by the applied FTSTS protocol led to the emergence of full network seizures, which continued spontaneously throughout the simulation. Figure 11D shows the decrease in the average E-to-I synaptic weight of the neocortical network induced by the applied FTSTS protocol. As shown here, the average E-to-I synaptic weight decreased by 30 nS immediately after the FTSTS protocol. Figures 11E and 11F show the percentage change in the synaptic strength between the excitatory and inhibitory neurons by the optimal inverted-standard FTSTS protocol during the suppression and enhancement of the prevalence of the epileptic seizures, respectively.

### Extension of Electrical FTSTS Protocol to Optogenetics

From the perspective of testing our designed FTSTS protocol in animal experiments, one of the potential limitations is that our electrical FTSTS protocol, described in the previous sections, requires spatially separable targeted excitatory and inhibitory populations. Although we addressed this limitation in the previous section by showing how our protocol can be applied to partially inseparable E-I networks, it may still be a limiting factor in many brain regions including the neocortex. To address this issue and to enable use of the FTSTS protocol in more realistic and practically realizable scenarios, in this section, we extend our electrical FTSTS protocol to an optogenetic-based FTSTS protocol (see Section 4 for details).

To test the efficacy of the optogenetic stimulation in the *in silico* seizure model, we initiated a seizure with a seizure initiating input applied for 3 *sec* shown as the red-square in Figure 12B. Then, we applied the optogenetic FTSTS protocol at 10 *sec* for a 10 *sec* duration. The optogenetic FTSTS was less efficient than the electrical stimulation with the bi-phasic pulse trains so the protocol required a 10 *sec* stimulation duration to increase the average synaptic weight by at least 15 *nS*, which is shown in Figure 12C. After the optogenetic FTSTS, we did not observe any spontaneous seizures for the rest of the simulation (see Figure 12B). Therefore, the integration of optogenetic techniques into our FTSTS protocol was able to overcome the spatial constraints of the FTSTS protocol in the neocortex and to prevent the emergence of spontaneous seizures.

**Figure 12:**
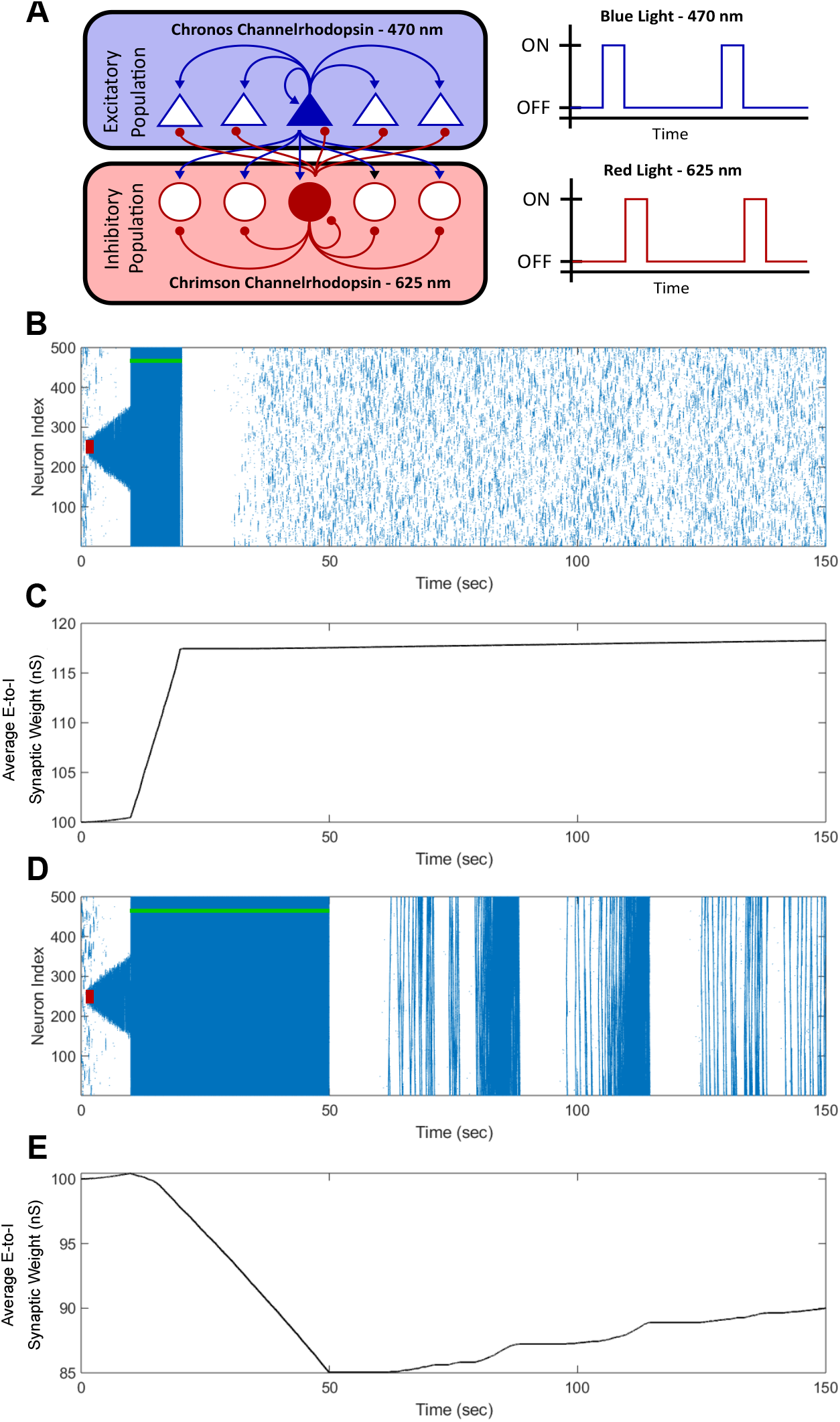
Optogenetic FTSTS Control of the Average Synaptic Weight. We incorporated optogenetic stimulation techniques into our FTSTS protocol to spatially and temporally selectively stimulate the excitatory and inhibitory neurons with blue (470 *nm*) and red (625 *nm*) light as shown in **(A)**. A seizure was initiated with a seizure initiating input of 200 *pA* applied for 3 *sec* shown as the red-box. Then, we applied our optical FTSTS to control the average synaptic weight of the network (green-bar). In order to increase the average synaptic weight, the Chronos channelrhodopsin was inserted into the excitatory neurons and the Chrimson channelrhodopsin was inserted into the inhibitory neurons. The raster plot in **(B)** shows the forced firing by the optical stimulation and that no further seizure were observed after the optogenetic FTSTS protocol. **(C)** shows the decrease in the average synaptic weight by the optogenetic FTSTS protocol. In order to decrease the average synaptic weight of the network, the Chronos channelrhodopsin was inserted into the inhibitory neurons and the Chrimson channelrhodopsin was inserted into the excitatory neurons. **(D)** shows the raster-plot of the excitatory neocortical neurons and the increased prevalence of strong seizures after the optogenetic FTSTS protocol. **(E)** shows the decrease in the average synaptic weight induced by the optogenetic FTSTS protocol.

**Figure 13:**
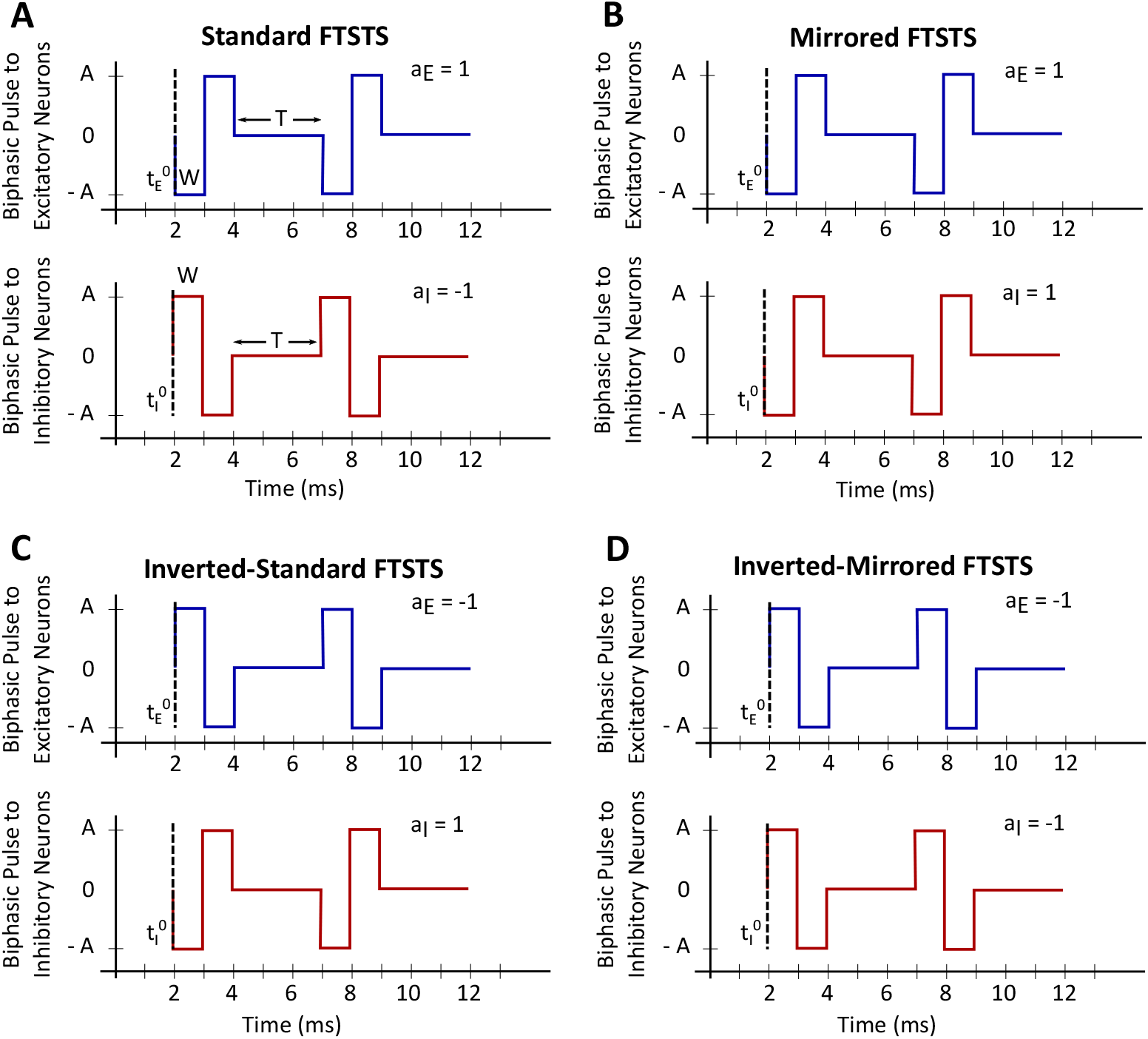
Forced Temporal Spike-Time Stimulation (FTSTS) parameters. The FTSTS pulse-pair parameters consist of amplitude (*A*), pulse width (*W*), pulse interval (*T*), and polarity (*a*_*E*_ and *a*_*I*_). The four different polarities are shown in **(A)** standard FTSTS, **(B)** mirrored FTSTS, **(C)** inverted-standard FTSTS, and **(D)** inverted-mirrored FTSTS.

After showing that the integration of the optogenetic technique into our FTSTS protocol was able to increase the average synaptic weight of the network, we examined whether the optogenetic FTSTS protocol would be able to decrease the average E-to-I synaptic weight of the network and increase the prevalence of spontaneous seizures, as well. Since the Chrimson channelrhodopsin dynamics were much slower than the Chronos channelrhodopsin dynamics, we swapped which channelrhodopsin was inserted into the excitatory and inhibitory neuron populations. In order to decrease the average synaptic strength, the Chronos channelrhodopsin was inserted into the inhibitory population and the Chrimson channelrhodopsin was inserted into the excitatory population. We were required to flip which channelrhodopsin was inserted into each population because the slower Chrimson dynamics caused unwanted neuron firing after optically stimulating the Chronos channelrhodopsin. Then, we initiated a seizure with a seizure initiating input (red-box) and applied the optogenetic FTSTS protocol at 10 *sec* for a 40 *sec* duration as shown in Figure 12D. The optogenetic FTSTS protocol stimulated the inhibitory neurons with blue (470 *nm*) light 2 *ms* before stimulating the excitatory neurons with red (625 *nm*) light. We applied the optogenetic FTSTS protocol for 40 *sec* to decrease the average synatpic weight 15 *nS* as shown in Figure 12E. After the FTSTS protocol, very strong seizures across the entire neocortical network emerged for the rest of the simulation. This highlights the ability of the optogenetic FTSTS to spatially and temporally selectively stimulates the excitatory and inhibitory neurons to not only increase the synaptic weight but also decrease the average synaptic weight of the network.

## 3 Discussion

In this paper, we systematically investigated the parameter space of our previously developed “Forced Temporal Spike-Time Stimulation” (FTSTS) strategy [1] to determine the efficacy of the FTSTS in reducing or increasing the neocortical-onset seizure prevalence. By harnessing the long-term synaptic plasticity between excitatory-to-inhibitory (E-to-I) connections in a *in silico* seizure model, we first showed that our FTSTS strategy (inverse-standard FTSTS) can effectively stop seizures and prevent further spontaneous seizures by increasing the average E-to-I synaptic weight (see Figure 2). By applying the exact reverse of our FTSTS protocol (standard FTSTS), we showed that the applied FTSTS protocol can increase the prevalence of spontaneous seizures by decreasing the average E-to-I synaptic weight (see Figure 3). Next, we explored the FTSTS pulse parameters, such as pulse amplitude, pulse-pair train-offset time, stimulation frequency, pulse width, and pulse polarity, on the efficacy of the FTSTS protocol in decreasing or increasing the prevalence of spontaneous seizures. Through our simulations, we found that the FTSTS efficacy enhanced monotonically with the pulse amplitude, the pulse width, and the inter-pulse interval (or stimulation frequency). Overall, we found that the most critical FTSTS parameters in determining the FTSTS efficacy were the train-offset time and the pulse-pair polarity. Tables 2 and 3 summarizes the best parameter combinations to increase and decrease the average E-to-I synaptic weight, respectively, from each of these investigations.

**Table 2:** List of best parameters to increase the average synaptic weight.

| Amplitude<br>(nA) | Pulse Width<br>(ms) | Frequency<br>(Hz) | Offset Time<br>(ms) | $a_E$ | $a_I$ | Rate of Weight Change<br>(nS/s) |
| --- | --- | --- | --- | --- | --- | --- |
| 2.0 | 2.0 | 83 | 3.0 | 1 | -1 | 5.281 |
| 2.0 | 2.0 | 83 | 2.0 | 1 | 1 | 5.397 |
| 2.0 | 1.0 | 83 | 0.5 | -1 | -1 | 6.161 |
| 2.0 | 1.0 | 83 | -0.5 | -1 | 1 | 6.237 |

**Table 3:** List of best parameters to decrease the average synaptic weight.

| Amplitude<br>(nA) | Pulse Width<br>(ms) | Frequency<br>(Hz) | Offset Time<br>(ms) | $a_E$ | $a_I$ | Rate of Weight Change<br>(nS/s) |
| --- | --- | --- | --- | --- | --- | --- |
| 2.0 | 2.0 | 83 | 2.0 | 1 | -1 | -4.373 |
| 2.0 | 2.0 | 83 | -0.5 | 1 | 1 | -4.77 |
| 2.0 | 1.0 | 83 | -0.5 | -1 | -1 | -5.589 |
| 2.0 | 1.0 | 83 | -2.0 | -1 | 1 | -5.933 |

Throughout our investigation, we found that the train-offset time induced the largest changes in the E-to-I synaptic weights. As the train-offset time between the two pulses delivered to the excitatory and inhibitory neuron populations was shifted, the efficacy of the FTSTS protocol in terminating or facilitating seizure activity drastically changed. For example, as shown in Figure 7A, when the train-offset time was changed from 0.5ms to 2ms, the effect of the FTSTS protocol flipped from decreasing to increasing the excitatory-to-inhibitory average synaptic weight, or equivalently from increasing to reducing the prevalence of spontaneous seizures. Since the FTSTS protocol is based on controlling the spike-times of the two neuron populations, it follows that slightly shifting the offset time between the two pulse trains will have a significant effect on the efficacy of the stimulation protocol. This result has a broader implication to already existing desynchronizing stimulation protocols, such as, coordinated reset [12, 13] and deep brain stimulation (DBS) [10, 11, 14]. While it was shown in [1] that integration of the FTSTS protocol into coordinate reset could improve the desynchronization protocol, these results showed that offsetting stimulation trains of the different stimulating electrodes may further improve the efficacy of the desynchronization protocol by forcing spiking patterns in different neural population that could result in the long-term desynchronization of a synchronous neural network. Additionally, if multiple high frequency stimulation (HFS) stimulating electrodes are inserted into separate neural populations for DBS, then offsetting the pulse trains of each electrode may improve the existing FDA approved neurostimulation-based therapies for brain disorders, such as epilepsy [11] and Parkinson’s Disease [14].

Our investigation into the effect of the inter-pulse interval (stimulation frequency) on the efficacy of the FTSTS protocol highlighted that the smaller inter-pulse interval (equivalently, higher stimulation frequency) is more effective in suppressing or enhancing the prevalence of spontaneous seizures. One exception, we observed, to this trend occurred when the inter-pulse interval was decreased from 10ms (83Hz) to 5ms (142Hz). Here, the efficacy generally decreased, since the lower inter-pulse intervals (higher stimulation frequencies) produced an undesired spike-time pattern that counteracted the desired spike-time temporal pattern. While the optimal FTSTS protocol inter-pulse interval was 10ms, large inter-pulse intervals (lower frequencies) were still able to control the E-to-I synaptic weights. These lower frequencies might be more desirable in experiments or potentially in neurostimulation devices for epileptic patients, since the lower frequency may lead to fewer side effects. Furthermore, if our protocol would be used to stop and prevent spontaneous seizures in *in vivo* experiments or in epileptic patients, the inter-pulse interval (stimulation frequency) could be used in a closed-loop control protocol to oppose quickly or slowly the changes in synaptic weights mediated by higher or lower frequencies, respectively.

In general, we found that the pulse amplitude and the pulse width influenced the efficacy of the FTSTS protocol as expected. An increase in the pulse amplitude increased the efficacy of the FTSTS protocol, but the efficacy plateaued when the amplitude increased beyond 2nA. Additionally, we observed that smaller pulse widths favored a more effective FTSTS protocol. This results from the higher temporal specificity in the induced spike-times of each neuron population, which reduces extra unwanted induced spiking. Importantly, we showed that even with the extra spiking induced by longer pulse width, the FTSTS protocol was able to control the average E-to-I synaptic weight of the network. This is critical for implementation of the FTSTS strategy into animal experiments or clinical trials where the stimulation pulse width may have a lower width limit.

In our electrical-based FTSTS protocol, we have assumed that the excitatory and inhibitory populations are well separated. This might not be the case for many brain regions including neocortex. In order to address this limitation, we integrated optogenetic stimulation dynamics into our electrical-based FTSTS protocol. Particularly, we inserted the Chronos channelrhodopsin into the excitatory population and the Chrimson channelrhodopsin into the inhibitory population. One of the advantages of optogenetic based stimulation is that it has high spatial selectivity. Since the FTSTS protocol requires the selective stimulation of separate neuron populations, optogenetic stimulation would be ideal for implementing FTSTS in animal experiments. Our simulation results showed that our optogenetic FTSTS protocol could effectively control the E-to-I synaptic weights in the neocortical-onset seizure network model (see Figure 12). One potential experiment that would confirm the efficacy of our stimulation protocol would be to optogenetically stimulate the excitatory neurons in layers 1-3 of the cortex and the inhibitory neurons in layers 2/3 [15] and record the changes in the dynamics of the excitatory and inhibitory neurons population in response to the optogenetic FTSTS. If the FTSTS increases the average synaptic weight between the excitatory and inhibitory neocortical neurons, and the inhibitory population activity increases, this would suggest that the FTSTS protocol is able to control the synaptic strength between the two populations.

In this work, our goal was to investigate the parameter space of the FTSTS protocol to determine the efficacy of the FTSTS protocol in controlling the excitatory-to-inhibitory synaptic weights and as a consequence, the seizure state of the biophysical neocortical-onset seizure model. While we determined the optimal FTSTS parameters to control the synaptic weights, our protocol relies on the assumption that the excitatory and inhibitory neurons can be separately stimulated. We show that this major limitation could be overcome by integrating the FTSTS protocol with an optogenetic stimulation protocol of the two neuron populations. Additionally, the *in silico* seizure model we used [2] only considered neocortical excitatory and inhibitory neurons and ignored any interconnectivity between neocortical neurons and neurons from other brain regions, such as the thalamus [15, 16]. While we didn’t consider internetwork connections with neurons of other brain regions, our protocol could be easily tested for this scenario. Also, the more separable the neuron population are from each other, more the efficacy and practicality of the FTSTS protocol would increase. Throughout this work, we only considered an open-loop FTSTS protocol. One future direction of this project would be to close-the-loop and develop an optimal closed-loop framework to prevent the emergence of seizures to treat epilepsy. Finally, one of the ultimate goals of this project is to show in experiment that the FTSTS protocol is able to control the synaptic weight between two neuron populations and control the synchronous state of a pathological synchronous neuron network.

## 4 Materials and methods

### Neocortical Seizure Model

Throughout this paper, we have used a recently published computational model of neocortical-onset seizures [2], validated with clinical data from epileptic patients, to simulate the seizure dynamics. Briefly, the model consists of a spatially homogeneous one-dimensional neural network consisting of 500 excitatory and 500 inhibitory neurons. The membrane potential, *V*, of each neuron is described by the following conductance-based integrate-and-fire model:

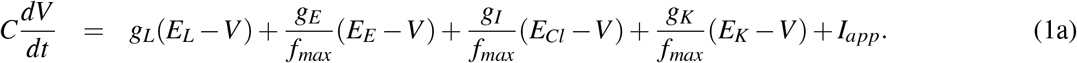

Here, *C* is the membrane capacitance, and *I*_*app*_ is the external electrical current. The model considers four conductances: leaky (*g*_*L*_), glutamatergic syanptic (*g*_*E*_), GABAergic synaptic (*g*_*I*_), and slow after hyperpolarization sAHP (*g*_*K*_). *f*_*max*_ is a scaling term equal to the maximum firing rate or the inverse of the refractory period. Each conductance has a corresponding reversal potential *E*_*L*_, *E*_*E*_, *E*_*I*_, and *E*_*K*_. The spike-times are stochastically determined based on the instantaneous firing rate 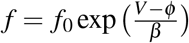, where *f*_0_ is a scaling parameter, *ϕ* is firing threshold, and *β* is the uncertainty of an action potential threshold. If a spike occurs, the membrane potential is set to average of 40 mV and the current membrane potential *V* (*t*_*i*_) at *t*_*i*_, where *t*_*i*_ is the time of a spike. After the spike, the membrane potential is reset to *V* (*t*_*i*+1_) = *V* (*t*_*i*_) − 20 mV at *t*_*i*+1_. The dynamics of the firing threshold *ϕ* that dictates the probability of a spike occurring is governed by Eq. (1b).

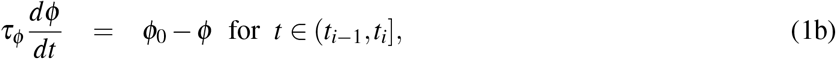

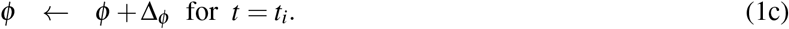

Here, *τ*_*ϕ*_ is a time constant, *ϕ*_0_ is the baseline threshold. After the spike at *t* = *t*_*i*_, *ϕ* is reset to *ϕ* (*t*_*i*_) + Δ_*ϕ*_. The excitatory and inhibitory synaptic dynamics are captured by *g*_*E*_ and *g*_*I*_, respectively (see Eq. (1a)). The conductance dynamics of the excitatory and inhibitory synapses are governed by the exponential decay functions shown in Eqs. (1d) and (1e).

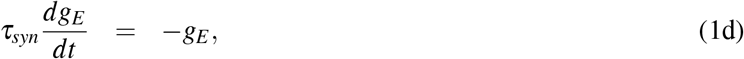

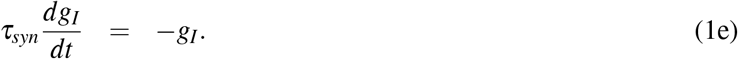

Here, *τ*_*syn*_ is the time constant. If an excitatory or inhibitory neuron spikes, then the weight of the synapse between the *i*^*th*^ pre-synaptic neuron and the *j*^*th*^ post-synaptic neuron is added to the *j*^*th*^ post-synaptic neuron’s conductance (*i*.*e*., *g*_*j*_(*t*_*i*_) = *g*_*j*_(*t*_*i*−1_) + *W* (*i, j*)). Furthermore, the inhibitory GABAergic synaptic input reversal potential is dependent on the chloride concentration gradient. The gradient determines the reversal potential (*E*_*Cl*_) of the GABAergic synapse using the following Nernst equation:

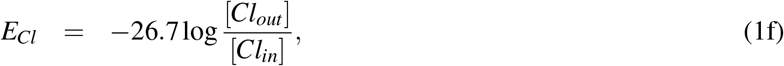

where [*Cl*_*out*_] and [*Cl*_*in*_] are the external and internal chloride concentration, respectively. The external chloride concentration is assumed to be constant and the internal chloride dynamics is modeled using the following equation:

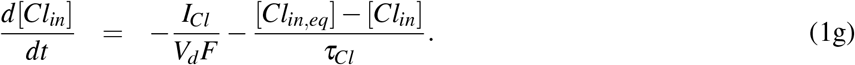

Here, the chloride current *I*_*Cl*_ is defined by *I*_*Cl*_ = *g*_*I*_(*V* − *E*_*Cl*_), *V*_*d*_ is the volume of distribution of [*Cl*_*in*_], *F* is the Faraday constant, and *τ*_*Cl*_ is the time constant. The equilibrium intracellular concentration of chloride is [*Cl*_*in,eq*_]. Finally, the slow after-hyperpolarization (sAHP) conductance dynamics *g*_*K*_ is modeled as

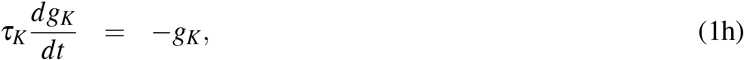

where *τ*_*K*_ is a time constant. If a spike occurs, then 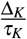 is added to *g*_*K*_ just after the spike. We provide the parameters for Eqs. (1a) - (1h) in Table 4.

**Table 4:** Model parameters of spiking neurons [2]

| Neuron Parameters | Value | Neuron Parameters | Value |
| --- | --- | --- | --- |
| $C$ | 100 pF | $\Delta\phi$ | -55 mV |
| $g_L$ | 4 nS | $\tau_{Cl}$ | 5000 mS |
| $E_L$ | -57 mV | $V_d$ | 0.2357 pL |
| $E_E$ | 0 mV | $[Cl]_{in,eq}$ | 6 mM |
| $E_K$ | -90 mV | $[Cl]_{out}$ | 110 mM |
| $f_0$ | 0.002 Hz | $\tau_K$ | 5000 ms |
| $\beta$ | 1.5 mV | $\Delta_K$ | 40 nS |
| $\tau_{ref}$ | 5 ms | $\sigma_E$ | 0.02 |
| $\tau_{syn}$ | 15 ms | $\sigma_I$ | 0.03 |
| $\tau_\phi$ | 100 ms | $f_{max}$ | 0.2 Hz |
| $\phi_0$ | -55 mV | $F$ | 96500 C mol<sup>-1</sup> |
| $\eta$ | 10<sup>-3</sup> | $\tau_{STD P}$ | 15 ms |

### Network Synaptic Connectivity

The synaptic connection between an excitatory and an inhibitory neuron is modeled in a distance-dependent form, as shown in Eqs. (2a) and (2b) [2].

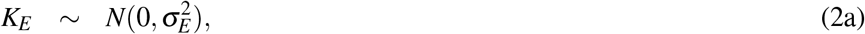

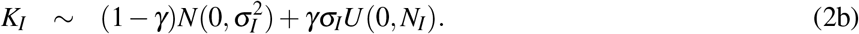

Briefly, the strength of the excitatory synapses is normally distributed about each excitatory neuron with mean zero and spatial variance of 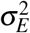, as described by the spatial distribution kernel *K*_*E*_ in Eq. (2a). The strength of inhibitory synaptic connections is modeled by the spatial distribution kernel *K*_*I*_ (see Eq. (2b)). In addition to the normally distributed synaptic strength with mean zero and spatial variance of 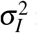 in Eq. (2b), there also exists weak uniformly distributed synapses in the inhibitory network, which are represented by the term *U*. The strength of the weak uniformly distributed synapses is *γσ*_*I*_. The parameter *γ* determines the contributions of the normally distributed and uniformly distributed synapses. The excitatory synaptic spatial kernel (*K*_*E*_) is used to define the initial synaptic weight matrix from the excitatory-to-inhibitory (*W*^*EI*^) and excitatory-to-excitatory (*W*^*EE*^) synapses. The inhibitory synaptic spatial kernel (*K*_*I*_) is used to define the initial synaptic weight matrix from the inhibitory-to-excitatory (*W*^*IE*^) and inhibitory-to-inhibitory synapses (*W*^*II*^).

### Spike-Time Dependent Plasticity Model

Throughout this work, we have assumed that E-to-E and E-to-I synapses in the neocortical model are plastic [2]. Furthermore, we have assumed that the change in the synaptic strength is activity-dependent, and we modeled the changes in the synaptic strength of both type of synapses using a Hebbian-based spike-timing dependent plasticity (STDP) rule [17, 18]. Eqs. (3a)-(3e) describe the STDP dynamics.

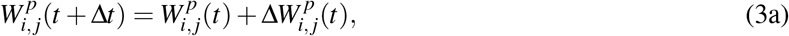

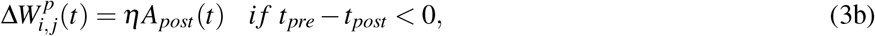

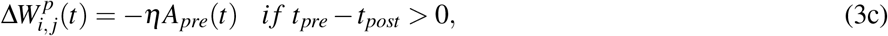

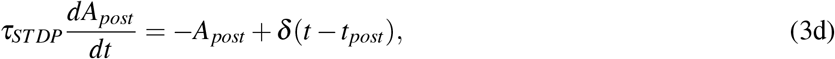

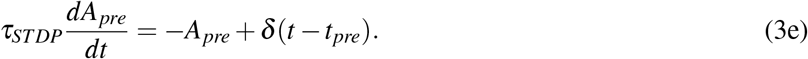

Here, 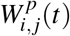 represents the fractional change in the weight (or strength) of a synapse connecting the *j*^*th*^ presynaptic neuron to the *i*^*th*^ postsynaptic neuron. The synaptic weight fraction at time *t* + Δ*t* (i.e., 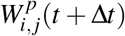) is updated by the change in the synaptic weight 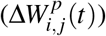, where Δ*t* is the time difference between the *j*^*th*^ presynaptic (*t*_*pre*_) and *i*^*th*^ postsynaptic (*t*_*post*_) neuron spike times, i.e., Δ*t* = *t*_*pre*_ −*t*_*post*_ (see Eq. (3a)). The synaptic weight of the synapse connecting the *j*^*th*^ presynaptic neuron to the *i*^*th*^ postsynaptic neuron is determined by 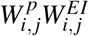 or 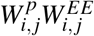 where the synaptic weight matrices *W*^*EI*^ and *W*^*EE*^ are derived from the excitatory spatial distribution kernal described in Eq. (2a). The parameter *η* represents the learning rate, i.e., the rate at which the synaptic weight is updated each time. The contributions of the long-term potentiation (LTP) and long-term depression (LTD) to the overall change in the synaptic weight, depending on the time difference between postsynaptic and presynaptic neuron spike-times, are modeled using the exponential functions *A*_*post*_(*t*) and *A*_*pre*_(*t*), respectively (see Eqs. (3d) and (3e)). The STDP time constant *τ*_*STDP*_ defines the the STDP spike time window [17]. At the time of the postsynapitc neuron spiking (*t*_*post*_) or the presynaptic neuron spiking at (*t*_*pre*_), Eqs. (3d) and (3e) are updated by the dirac-delta functions *δ* (*t* −*t*_*post*_) = 1 and *δ* (*t* −*t*_*pre*_) = 1, respectively. We provide the STDP parameters used in this paper in Table 4.

### Forced Temporal Spike-Timing Stimulation

In this work, we will use our previously developed Forced Temporal Spike-Timing Stimulation (FTSTS) strategy [1], which has shown to be effective in controlling the synchronization of neurons in E-I networks by harnessing the synaptic plasticity of the network. Briefly, our FTSTS protocol consists of excitatory and inhibitory charge-balanced biphasic stimulation pulses delivered to individual neurons in each of the subpopulation. The positive portion of each individual charge-balanced biphasic pulse of the FTSTS protocol forces the neurons to fire, while the negative portion of the biphasic pulse prevents neurons from firing. An example of the standard FTSTS protocol is shown in Figure 13A. We showed in [1] that forcing the neurons in the excitatory and inhibitory populations to fire in a postsynaptic (inhibitory) neuron before presynaptic (excitatory) neuron firing pattern decreased the excitatory-to-inhibitory synaptic weight. Additionally, if we forced a presynaptic (excitatory) neuron to fire an action potential before the postsynaptic (inhibitory) neuron, it increased the excitatory-to-inhibitory synaptic weight. In order to induce one of these firing patterns to increase or decrease the synaptic weight, separate biphasic pulse trains were applied to the excitatory and inhibiotory neuron populations such that the biphasic pulse train applied to the excitatory population was the exact inverse of the biphasic pulse train applied to the inhibitory population. For example, if the positive portion of the biphasic pulse is applied to the excitatory neuron population, then the negative portion of the biphasic pulse is applied to the inhibitory neuron population, and *vice versa* when the negative portion of the biphasice pulse is applied to the excitatory neuron population. To induce a postsynaptic before presynaptic firing pattern to decrease the synaptic weight that we call standard FTSTS, the positive portion of the biphasic pulse is first applied to the inhibitory population while the negative portion of the biphasic pulse is applied to the excitatory population. Then, the negative portion of the biphasic pulse to the inhibitory is applied while the positive portion of the biphasic pulse is applied to the excitatory neurons.

**Figure 14:**
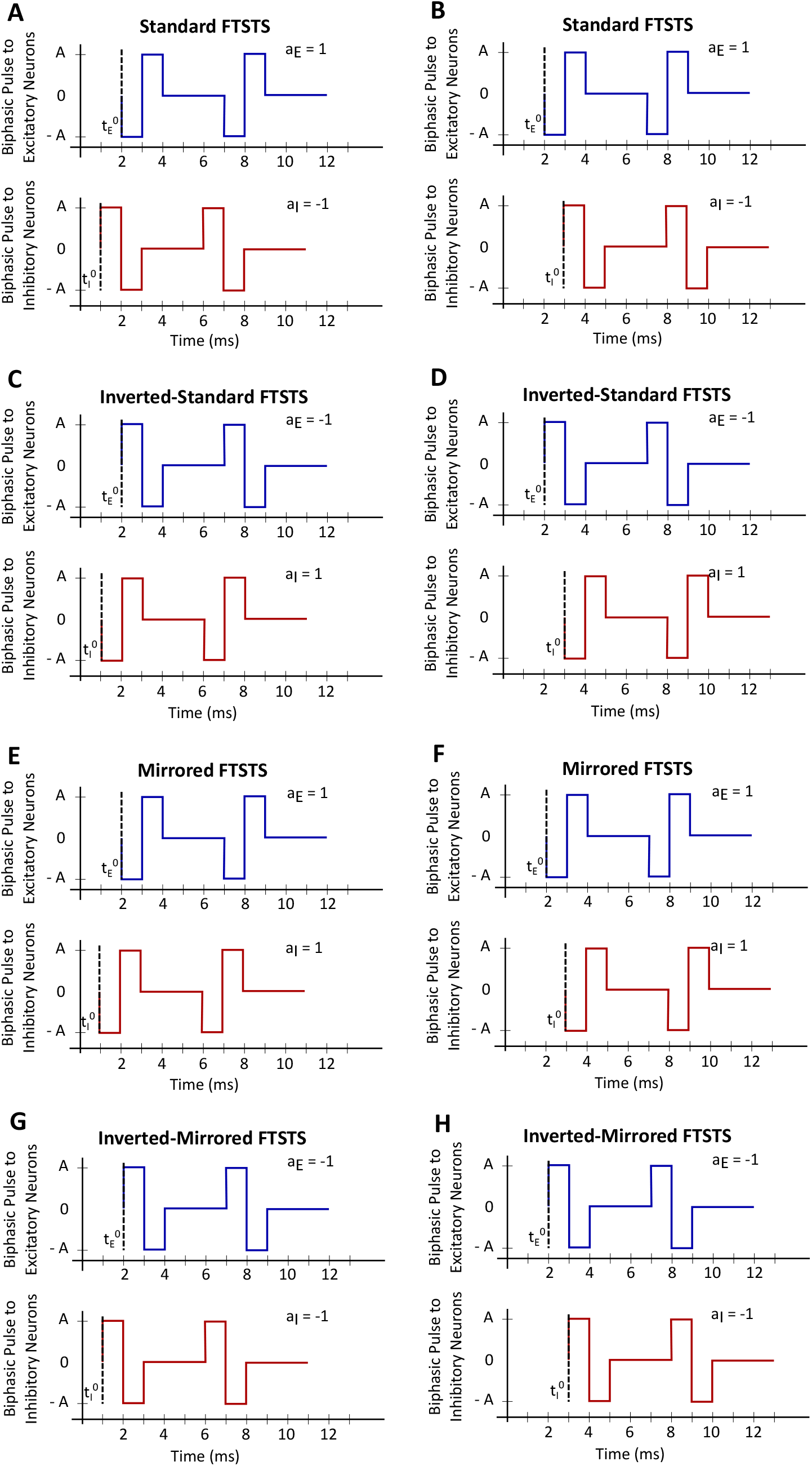
The train-time offset of the FTSTS pulse-pair. **(A), (C), (E)** and **(G)** show examples of a train-offset time of −1 *ms* for standard FTSTS, inverted-standard FTSTS, mirrored FTSTS, and inverted-mirrored FTSTS, respectively. **(B), (D), (F)** and **(H)** show examples of a train-offset time of 1 *ms* for standard FTSTS, inverted-standard FTSTS, mirrored FTSTS, and inverted-mirrored FTSTS, respectively.

In order to fully consider the parameter space of the FTSTS protocol in this work, we have modified our previously developed FTSTS protocol as follows. Particularly, we have defined a FTSTS biphasic pulse based on its polarity. We say that the FTSTS pulse has a polarity of +1 if the biphasic pulse begins with the negative amplitude. Similarly, if a biphasic pulse begins with the positive amplitude, we say that it has a polarity of −1. Based on this definition, we have defined four FTSTS protocols: (1) the standard FTSTS in which the neurons in the excitatory population receive a biphasic stimulation pulse with polarity of +1 (i.e., *a*_*E*_ = +1) and the neurons in the inhibitory population receive a biphasic stimulation pulse with polarity of −1 (i.e., *a*_*I*_ = −1); (2) the inverted-standard FTSTS protocol in which the neurons in the excitatory population receive a biphasic stimulation pulse with polarity of −1 (i.e., *a*_*E*_ = −1) and the neurons in the inhibitory population receive a biphasic stimulation pulse with polarity of +1 (i.e., *a*_*I*_ = +1) ; (3) the mirrored FTSTS protocol in which the neurons in both the excitatory and inhibitory populations receive a biphasic stimulation pulse with polarity of +1 (i.e., *a*_*E*_ = *a*_*I*_ = +1) ; and (4) the inverted-mirrored FTSTS protocol in which the neurons in both the excitatory and inhibitory populations receive a biphasic stimulation pulse with polarity of −1 (i.e., *a*_*E*_ = *a*_*I*_ = −1). In addition to the FTSTS pulse polarity as a parameter, we have also considered the pulse amplitude, pulse width, pulse interval (or the stimulation frequency), and pulse-pair train-offset time (Δ*ϕ*, which we defined as the time difference between the start time of the inhibitory population pulse 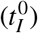 and the start time of the excitatory population pulse 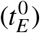 as shown in Eq. (4)) as the parameters of the FTSTS protocol to be optimized. Figure 13A illustrates these parameters.

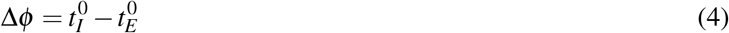

### Optogenetic FTSTS Protocol

To incorporate an optogenetic-based stimulation in our FTSTS protocol, we modified the dynamics of the neocortical-seizure model by incorporating two different optogenetic channelrhodopsin ion channels, namely, the channelrhodopsin Chronos into the excitatory neurons, which selectively responds to blue light (460 nm), and the channelrhodopsin Chrimson into the inhibitory neurons, which selectively responds to red light (625 *nm*) [19]. We modeled the channelrhodopsin light activated current using a reduced model developed in [20] and [21]. The channelrhodopsin dynamics induced by the optical stimulation are shown in Eqs. (5a) - (5h). The channelrhodopsin follows a simple exponential decay once the optical input is turned off. In order to incoporate the optogenetic dynamics into the neocortical-seizure model, the channelrhodopsin current (*I*_*ChR*2_) (see Eq. (5a)) is added to Eq. (1a). Eq. (5b) defines the conductance waveform *F*_*ChR*2_, which is dependent on both the intensity of the applied light (*W*_*light*_) and the duration the light is applied (*t* −*t*_*on*_ − *d*). Here, *t*_*on*_ is the time the optical stimulation is applied and *d* is a light dependent activation delay. *τ*_*act*_ and *τ*_*inact*_ represent the activation and deactivation time constants. The parameters of the light dependent delay variable *d* are *d*_*A*_, *d*_*B*_, and *d*_*C*_ (see Eq. (5c)). The light intensity dependent variables 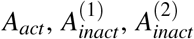, and *A*_*persist*_ modulate the open state of the channelrhodopsin. *A*_*act*_ represents the light dependent activation of the channelrhodopsin and is modeled using Eq. (5e). 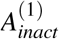 and 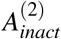 determine closing the rate of the channelrhodopsin and are modeled using Eqs. (5f)-(5g). Finally, *A*_*persist*_ is the persistent activation of the channelrhodopsin during a prolonged optical stimulation is modeled using Eq. (5h). The activation and deactivation parameters in Eqs.(5e)-(5g) are *a*_0_, *a*_*min*_, *b*_0_, *b*_1_, *b*_2_, *c*_*inact*_, and *k*_*inact*_.

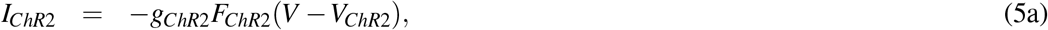

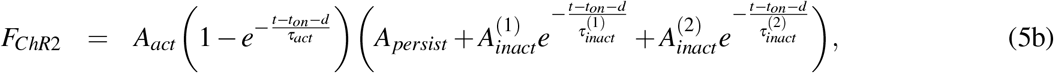

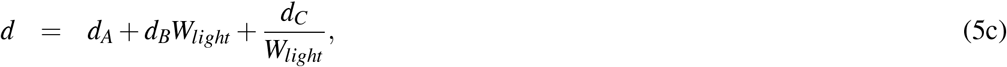

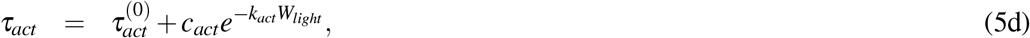

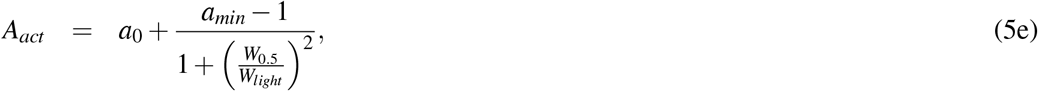

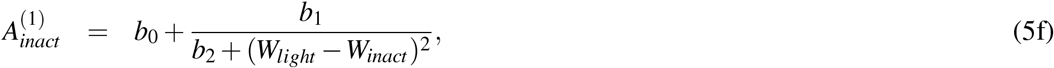

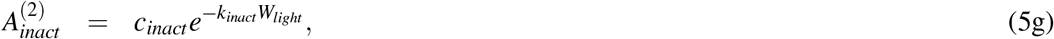

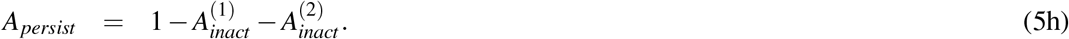

The exponential decay parameters were set as the experimentally measured values for each channelrhodopsin [19].

Additionally, the light needed to stimulate the two different channelrhodopsin ion channels (i.e., 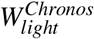 and 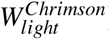) was determined from the literature, such that there was no light interference [19]. The channelrhodopsin parameters are listed in Table 5.

**Table 5:** Channelrhodopsin photocurrent parameters.

| Neuron Parameters | Value | Neuron Parameters | Value |
| --- | --- | --- | --- |
| $W_{inact}$ | 0.11 | $W_{light}^{Chronos}$ | 0.0308 |
| $\tau_{inact}^{(1)}$ | 9.06 ms | $W_{light}^{Chrimson}$ | 0.0023 |
| $d_A$ | 0.27 | $b_0$ | 0.16 |
| $d_B$ | -0.05 | $b_1$ | 0.013 |
| $d_C$ | -0.0126 | $b_2$ | 0.027 |
| $\tau_{act}^{(0)}$ | 0.74 ms | $\tau_{inact}^{(2)}$ | 59.6 ms |
| $c_{act}$ | 12 | $c_{inact}$ | 0.29 |
| $k_{act}$ | 25 | $k_{inact}$ | 2.4 |
| $a_0$ | 1 | $V_{ChR2}$ | 0 mV |
| $a_{min}$ | 0.4 | $g_{ChR2}$ | 294 nS |
| $W_{0.5}$ | 0.38 | $\tau_{off}^{Chronos}$ | 3.6 ms |
| | | $\tau_{off}^{Chrimson}$ | 15.8 ms |

## 5 Acknowledgments

We gratefully acknowledge funding and support from the San José State University and the University of Idaho Department of Chemical and Biological Engineering.

